# Modeling of DNA methylation in cis reveals principles of chromatin folding *in vivo* in the absence of crosslinking and ligation

**DOI:** 10.1101/407031

**Authors:** Josef Redolfi, Yinxiu Zhan, Christian Valdes, Mariya Kryzhanovska, Isabel Misteli Guerreiro, Vytautas Iesmantavicius, Guido Tiana, Tim Pollex, Jop Kind, Sebastien Smallwood, Wouter de Laat, Luca Giorgetti

## Abstract

Mammalian chromosomes are folded into an intricate hierarchy of structural domains, within which topologically associating domains (TADs) and CTCF-associated loops partition the physical interactions between regulatory sequences. Current understanding of chromosome folding largely relies on chromosome conformation capture (3C)-based experiments, where chromosomal interactions are detected as ligation products after crosslinking of chromatin. To measure chromosome structure *in vivo*, quantitatively and without relying on crosslinking and ligation, we have implemented a new method named damC. DamC combines DNA-methylation based detection of chromosomal interactions with next-generation sequencing and a biophysical model of methylation kinetics. DamC performed in mouse embryonic stem cells provides the first *in vivo* validation of the existence of TADs and CTCF loops, confirms 3C-based measurements of the scaling of contact probabilities within TADs, and provides evidence that mammalian chromatin *in vivo* is essentially rigid below 5 kilobases. Combining damC with transposon-mediated genomic engineering shows that new loops can be formed between ectopically introduced and endogenous CTCF sites, which alters the partitioning of physical interactions within TADs. This establishes damC as a crosslinking-and ligation-free framework to measure and modify chromosome interactions combined with a solid theoretical background for rigorous data interpretation. This orthogonal approach to 3C validates the existence of key structural features of mammalian chromosomes and provides novel insights into how chromosome structure within TADs can be manipulated.

## Introduction

Characterizing how chromosomes are folded is fundamental to better understand the control of gene expression and how it possibly constrains genome evolution. Chromosome conformation capture (3C) methods, and notably its derivatives relying on high-throughput sequencing such as 5C, Hi-C and 4C^1^, have significantly contributed to our current understanding of genome architecture. 3C-based methods have revealed that chromosome folding is driven by at least two independent mechanisms. On the one hand, the preferential (and mutually exclusive) associations between transcriptionally active or inactive chromosomal locations give rise to the so-called A and B compartments^2^. On the other hand, chromatin loops are formed between promoters and enhancers and between convergent CTCF binding sites, the latter through cooperative action between cohesin and the DNA-binding protein CTCF^3^. The interplay between compartmentalization and CTCF/cohesin-driven loop formation results in complex nested hierarchies of folding domains^4,5^, among which topologically associating domains (TADs)^6–8^ stand out as preferential functional units^9^. The involvement of CTCF in loop formation has been demonstrated using global depletion experiments^10,11^, as well as targeted deletions/inversions of CTCF sites leading to loss of looping interactions^12–14^. The underlying molecular mechanisms are however still incompletely understood. According to a highly influential hypothesis and supported by recent in vitro evidence^15^, CTCF-mediated interactions occur as cohesin extrudes chromatin loops until it is blocked by CTCF bound to DNA in a defined orientation^16^. According to this hypothesis, ectopic insertion of CTCF sites could result in the establishment of new loops onto endogenous CTCF sites, depending on their mutual orientation. Remarkably however, this has not been experimentally demonstrated.

In 3C, detection of spatial proximity is based on formaldehyde crosslinking followed by digestion and ligation of crosslinked chromatin^1^. Both crosslinking and ligation represent sources of potential experimental bias, raising the question as to whether the structures detected by 3C methods actually exist in living cells^17–20^. Importantly, 3C-based crosslinking frequencies are usually assumed to be proportional to absolute chromosomal contact probabilities and used to build mechanistic physical models of chromosome folding^21,22^, including the loop-extrusion model^16^. However there is no formal proof that 3C counts are proportional to actual chromosomal contact probabilities. Independent techniques such as DNA in situ hybridization (DNA FISH)^6,23^, genome architecture mapping (GAM)^24^, native 3C^25^ and split-pool recognition of interactions by tag extension (SPRITE)^26^ have also been used to detect loops, TAD boundaries and compartments. Nevertheless, these methods still involve substantial biochemical manipulation of cells, and employ either crosslinking or ligation. No crosslinking-and ligation-free method is currently available to detect chromosome interactions in living cells and remarkably, evidence that CTCF-associated loops actually exist is based exclusively on crosslinking methods.

To measure chromosomal contact probabilities *in vivo*, quantitatively and without relying on crosslinking and ligation, we have implemented a new method named ‘damC’ based on a modified version of DamID^27^ and a physical model of methylation kinetics. In damC, the DNA adenine methyltransferase Dam is inducibly recruited to specific genomic viewpoints, allowing the detection of chromosomal interactions with the binding site by high-throughput sequencing of methylated DNA across megabase-scale regions. Physical modeling of this process shows that the experimental output in damC is directly proportional to chromosomal contact probabilities, thus providing a theoretical framework for the interpretation of data.

Using damC we here provide the first crosslinking-and ligation-free validation of structures identified by 3C methods. Notably, by comparing damC with 4C-seq and high-resolution Hi-C at hundreds of genomic locations in mouse embryonic stem cells (mESC), we confirm the existence of TAD boundaries as well as CTCF-associated loops. We also show that the scaling of contact probabilities measured in damC is the same as in 4C and Hi-C, providing strong evidence in favor of current interpretations of 3C-based data in terms of physical models of chromosome folding. Using modified damC viewpoints containing CTCF binding sites, we additionally demonstrate that the ectopic insertion of CTCF sites can lead to the formation of new loops with endogenous sequences bound by CTCF. This shows that it is possible to manipulate local chromosome structure by inserting short ectopic sequences that rewire interactions within TADs.

## Results

### damC: Methylation-based detection of chromosomal contacts *in vivo*

In classical DamID, the bacterial DNA adenine methyltransferase Dam is fused to a DNA-binding protein of interest resulting in the ectopic methylation of adenines within GATC motifs in the immediate neighborhood of the protein DNA binding sites^27^. Methylated GATCs (GmATC) are then specifically digested by the DpnI restriction enzyme and detected using high-throughput sequencing. After normalisation for nonspecific methylation due to freely diffusing Dam molecules, this allows to determine the DNA binding location of the fusion protein. In principle, recruiting the methyltransferase to a specific genomic viewpoint should also allow to detect methylation at chromosomal regions that interact with the viewpoint in 3D, if interaction-specific methylation is appreciable over methylation by freely diffusing Dam. We reasoned that we could further explore this idea by fusing Dam to the reverse tetracycline receptor (rTetR), and inserting an array of Tet operators (TetO) in the genome. This would ensure the targeted, inducible recruitment of large numbers of Dam molecules to a specific genomic viewpoint in the presence of doxycycline (Dox) (**Figure 1a, left**), and maximize the chances that distal sites are methylated when interacting with the viewpoint. In the absence of Dox, rTetR-Dam does not bind to the viewpoint, allowing an accurate estimation of nonspecific methylation (**Figure 1a, right**) and precise background correction. Coupled to high-throughput sequencing, this strategy could provide 4C-like, ‘one vs. all’ interaction profiles^28^ measuring contact probabilities from the TetO viewpoint (**Figure 1a**) across large genomic distances and at high genomic resolution (one GATC site every ∼250bp on average). Insertion of multiple TetO arrays separated by large genomic distances would additionally allow the interrogation of chromosomal interactions in parallel from many viewpoints (**Figure 1b**). We refer to this method as ‘damC’.

**Figure 1.**
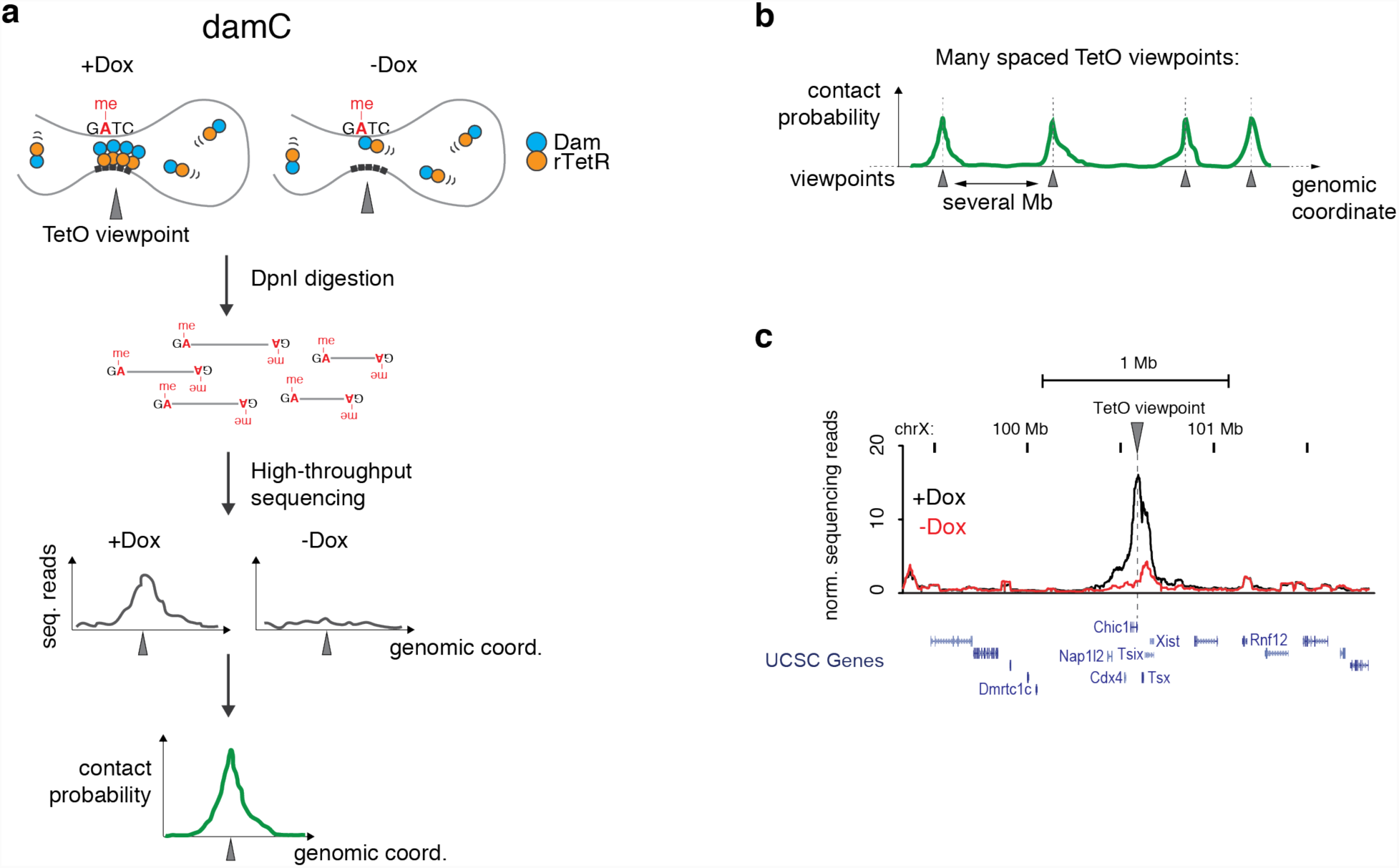
damC: methylation-based measurement of chromosomal interactions. **a)** Scheme of damC experiments. In the presence of doxycycline (+Dox) rTetR-Dam binds to a genomic viewpoint through a TetO array, and methylates adenines in GATC sites that contact the viewpoint. In the absence of doxycycline (-Dox) only nonspecific methylation by freely diffusing rTetR-Dam occurs. Methylated GATCs can be detected by digestion of genomic DNA with DpnI and next generation sequencing of the restriction sites. Correction for nonspecific methylation allows to extract contact probabilities with the TetO viewpoint. **b)** Insertion of multiple TetO arrays spaced by several Mb allows detecting interaction from single viewpoints in parallel. **c)** Proof-of-principle experiment showing increased methylation in *cis* following the recruitment of rTetR-Dam to an array of 256 TetO sites in the 5’UTR of the *Chic1* gene (black) in the presence of Dox, compared to the -Dpx control where rTetR-Dam is not recruited (red).

To test the experimental feasibility of this approach, we initially employed female mouse embryonic stem cells (mESCs) that carry one array of 256 TetO sites on chromosome X^29^. A subclone of these cells with only one copy of the X chromosome was transiently transfected with an expression plasmid for rTetR-Dam and methylation was measured after 24 hours^30^. This revealed significantly higher methylation upon induction with Dox compared to the uninduced control. Importantly this signal extended over approximately 300 kb around the TetO viewpoint (**Figure 1c**). This argued that targeted recruitment of Dam leads to increased methylation in *cis* over long genomic distances, consistent with previous observations using semi-quantitative schemes for the detection of methylation^25,31,32^. Importantly GmATCs can be easily recovered by high-throughput sequencing, allowing accurate measurement of methylation levels. Since methylation is determined by the interplay between the methyltransferase activity and passive demethylation during DNA replication, we reasoned that it should be possible to quantitatively model this process and derive chromosomal contact probabilities from the sequencing readout.

### The experimental output in damC is proportional to chromosomal contact probabilities

The methylation level of a single GmATC is determined by a dynamic interplay between methylation (by either freely diffusing or TetO-bound Dam) and demethylation due to DNA replication, when a fully methylated GmATC generates two hemi-methylated sites that are essentially not detected in DamID^33^ (**Figure 2a**). To identify an experimental quantity that would be directly proportional to chromosomal contact frequencies, we generated a physical model describing the time evolution of methylation at a GATC site located at an arbitrary genomic distance from the TetO viewpoint (**Supplementary Model description**). The model describes the time evolution of the number of GmATC sites in terms of rate equations (**Figure 2b**) which take into account that methylation by TetO-bound Dam only occurs in the presence of Dox (**Figure 1a**). Methylation rates are allowed to depend on local biases (e.g. chromatin accessibility or mappability). By solving the rate equations under the assumption that methylation is faster than demethylation (as observed in Ref.^33^) and the duration of the experiment, we found that the contact probability between the GATC site and the TetO viewpoint is directly proportional to a measurable quantity. This quantity, which we refer to as ‘damC enrichment’, is simply the relative difference between methylation levels in the presence and absence of Dox (**Figure 2c**). Physical modeling of methylation kinetics thus shows that damC can directly measure chromosomal contact probabilities.

**Figure 2.**
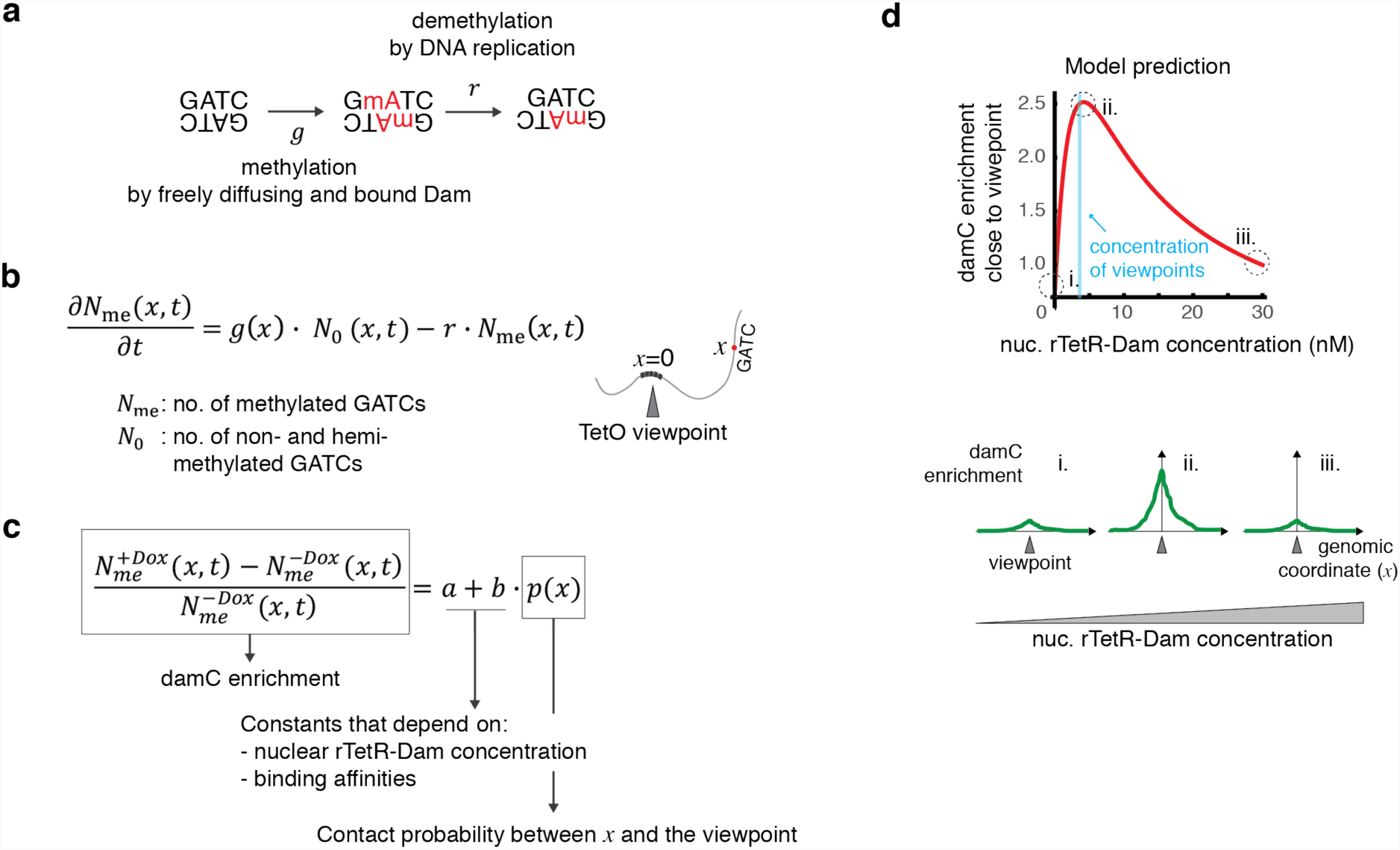
Physical model of methylation dynamics. **a)** Unmethylated GATCs can be methylated by either freely diffusing or TetO-bound rTetR-Dam with rate *g*, and partially demethylated during DNA replication occurring at rate *r*. Partially demethylated GATCs are inefficiently cut by DpnI and do not contribute to the damC experiment. **b)** Model of methylation dynamics. The time evolution of the number of methylated and non-/hemi-methylated GATCs located at genomic distance *x* from a TetO viewpoint is described in terms of ordinary differential equations governed by rates *g* and *r*. **c)** The damC enrichment at a generic location *x* is independent on time and proportional to the contact probability between *x* and the viewpoint. Proportionality constants *a* and *b* depend on the nuclear rTetR-Dam concentration and the binding affinities for the TetO and nonspecific genomic sites. **d)** Model prediction using example parameter values (rTetR-TetO affinity = 5 nM, nonspecific affinity = 80 nM, 600 TetO insertions corresponding to 2 nM in a nuclear volume of ∼490 fl, and contact probability of 0.5 corresponding to an interaction occurring in half of the cell population). The behavior of the curve is conserved across a wide range of physiologically relevant parameter values (see Supplementary Figure 1b).

Given a certain contact probability between the GATC site and the TetO viewpoint, the model predicts that the damC enrichment depends on three parameters: 1) the nuclear concentration of rTetR-Dam, 2) the rTetR-Dam binding affinity for the TetO array, and 3) the average nonspecific binding affinity of rTetR-Dam for endogenous genomic sites (**Figure 2c**). Importantly, the damC enrichment does not depend on local methylation biases and therefore should not be affected by differential accessibility or mappability, provided that interactions with the TetO viewpoint can increase methylation at the GATC site (i.e. local methylation is not saturated in the absence of Dox). In fact, once the binding affinities are fixed to certain values, the main determinant of damC enrichment in experimental conditions is the nuclear concentration of rTetR-Dam. In particular, damC enrichment is predicted to be maximal when rTetR-Dam concentration is around the nuclear concentration of TetO viewpoints (**Figure 2d** and **Supplementary Figure 1a**), and negligible when the rTetR-Dam concentration is either very high or very low. This behavior does not depend on the particular values of the affinity constants (**Supplementary Figure 1b**). Thus, physical modeling predicts that accurate control of rTetR-Dam nuclear concentrations is needed to perform damC with optimal signal-to-noise ratio.

### damC from hundreds of genomic viewpoints validates model predictions

To test the model predictions and experimentally measure chromosomal interactions using damC, we established a mESC line allowing the control of the nuclear concentration of rTetR-Dam. To this purpose we created a stable cell line expressing rTetR fused with enhanced green fluorescent protein (EGFP), Dam, and the mutant estrogen ligand-binding domain ERT2. ERT2 ensures the cytoplasmic localization of the fusion protein in the absence of 4-hydroxy-tamoxifen (4-OHT), thus preventing constitutive GATC methylation. It also enables to control its nuclear level by changing the 4-OHT concentration in the culture medium (**Figure 3a**) as confirmed by nuclear accumulation of the EGFP signal as a function of increasing 4-OHT dose (**Supplementary Figure 2a**).

**Figure 3.**
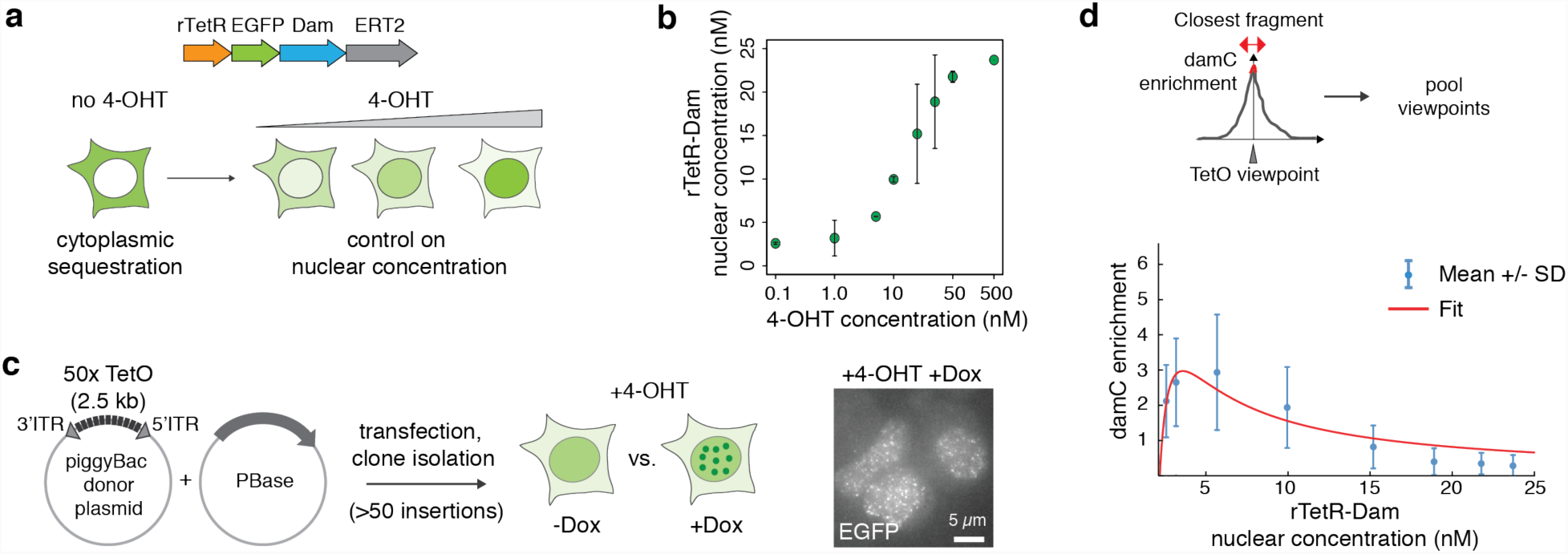
An inducible mESC line to perform damC and test the model predictions. **a)** mESCs expressing rTetR-Dam-EGFP-ERT2 allow to control the nuclear concentration of the fusion protein by changing the amount of 4-hydroxy-tamoxifen (4-OHT) in the culture medium. **b)** Nuclear concentration of the rTetR-Dam fusion protein as a function of 4-OHT concentration. Number of protein copies per nucleus were determined using mass spectrometry on nuclear extracts and divided by the average nuclear volume (∼490 fl) determined using DAPI staining (see Supplementary Figure 2). Error bars corresponds to the standard deviation of the two experimental replicates. **c)** Random integration of large numbers of 50x TetO platforms using the piggyBac transposon. Accumulation of EGFP signal to nuclear foci in the presence of Dox (right: max. intensity projection over 10 Z planes) indicates binding of rTetR-Dam to the arrays and allows selecting clones with large numbers of insertions. **d)** Quantification of damC experiments as a function of rTetR-Dam concentration. damC enrichment in the immediate vicinity of TetO viewpoints (contact probability ∼1) shows a maximum in line with the model prediction. Blue: Average +/- standard deviation over the 100 TetO viewpoints with highest enrichment. Red line: model fit to the experimental data.

To quantitatively measure the nuclear concentration of the rTetR-Dam fusion protein as a function of 4-OHT concentration, we analyzed nuclear protein extracts using mass spectrometry. Combining the proteomic ruler strategy^34^ with parallel reaction monitoring (PRM) (see **Methods, Supplementary Figure 2b-c** and **Supplementary Table S1**) we estimated the nuclear rTetR-Dam concentration to vary gradually between approximately 3 and 25 nM when increasing the 4-OHT concentration from 0.1 to 500 nM (**Figure 3b**).

In order to generate a large number of TetO viewpoints in the same cell line, we randomly inserted arrays of 50 TetO sites (each spanning approx. 2.7 kb) using the piggyBac transposon system^35^ (**Figure 3c**). This resulted in clonal mESC lines carrying at least 100 TetO arrays, judging from the accumulation of EGFP at nuclear foci in the presence of 4-OHT and Dox (**Figure 3c**). We further selected one clone carrying 890 TetO array insertions (**Supplementary Table S2**), as determined by high-throughput mapping of piggyBac insertion locations (**Methods** and **Supplementary Table S2**).

We then performed damC after treating cells overnight with different doses of 4-OHT in the presence and in the absence of Dox. Experiments were performed using a new next-generation sequencing library preparation protocol that includes unique molecular identifiers (UMI) and increases the coverage of methylated GATC sites genome-wide, thus maximizing the proportionality between methylation levels and sequencing readout (**Supplementary Figure 2d-e** and **Methods**). We quantified damC enrichment as in **Figure 2c** in the immediate vicinity of TetO viewpoints, and plotted it as a function of the rTetR-Dam concentration determined in mass spectrometry (**Figure 3d**). Consistent with the model prediction that damC enrichment should be maximal when the rTetR-Dam nuclear concentration is in the range of the nuclear concentration of TetO viewpoints, we found that the maximum occurs at 2.7 nM corresponding to 795 viewpoints per nucleus. Model fitting resulted in an estimate of 0.6 nM for the specific rTetR-TetO binding constant, in the range of values obtained in vitro^36^, and 18nM for the average non-specific binding constant accounting for rTetR and Dam interactions with GATC sites genome-wide.

These results provide a validation of the damC model and therefore support the interpretation of the damC enrichment in terms of contact probabilities. They additionally highlight that in our experimental system, the most significant amounts of damC enrichment can be observed in a range of rTetR-Dam nuclear concentrations corresponding to 5-10 nM 4-OHT (**Supplementary Figure 2f**).

### damC reveals the existence of TADs and loops *in vivo*

Under optimal 4-OHT concentrations (10 nM), zooming into individual TetO viewpoints revealed significant damC enrichment extending over hundreds of kilobases around each viewpoint (**Fig. 4a**). Since biological replicates were highly correlated (**Supplementary Figure 3a**), we analyzed the merged data. Importantly, damC enrichment profiles showed remarkable agreement with 4C experiments obtained using the same TetO arrays as viewpoints and DpnII as a primary restriction enzyme (**Figure 4a** and **Methods**). The damC enrichment was systematically concentrated within TAD boundaries detected in Hi-C experiments performed on the same cell line (**Figure 4a**) and steeply decayed across TAD boundaries by roughly a factor two, in excellent agreement with 4C (**Figure 4c**). Of the 100 insertions showing the highest signal-to-noise ratio (listed in **Supplementary Table S3**), only a handful occurred in close proximity (within 1kb) with either an active regulatory element, or a CTCF site (**Supplementary Figure 3b**). Also in these cases, the damC enrichment profiles were highly overlapping with 4C (**Figure 4b**), with both strategies revealing specific looping interactions between endogenous convergent CTCF sites in vivo (**Figure 4b, right**) which were confirmed using the partner CTCF sites as a reciprocal viewpoint in 4C (**Supplementary Figure 3c)**.

**Figure 4.**
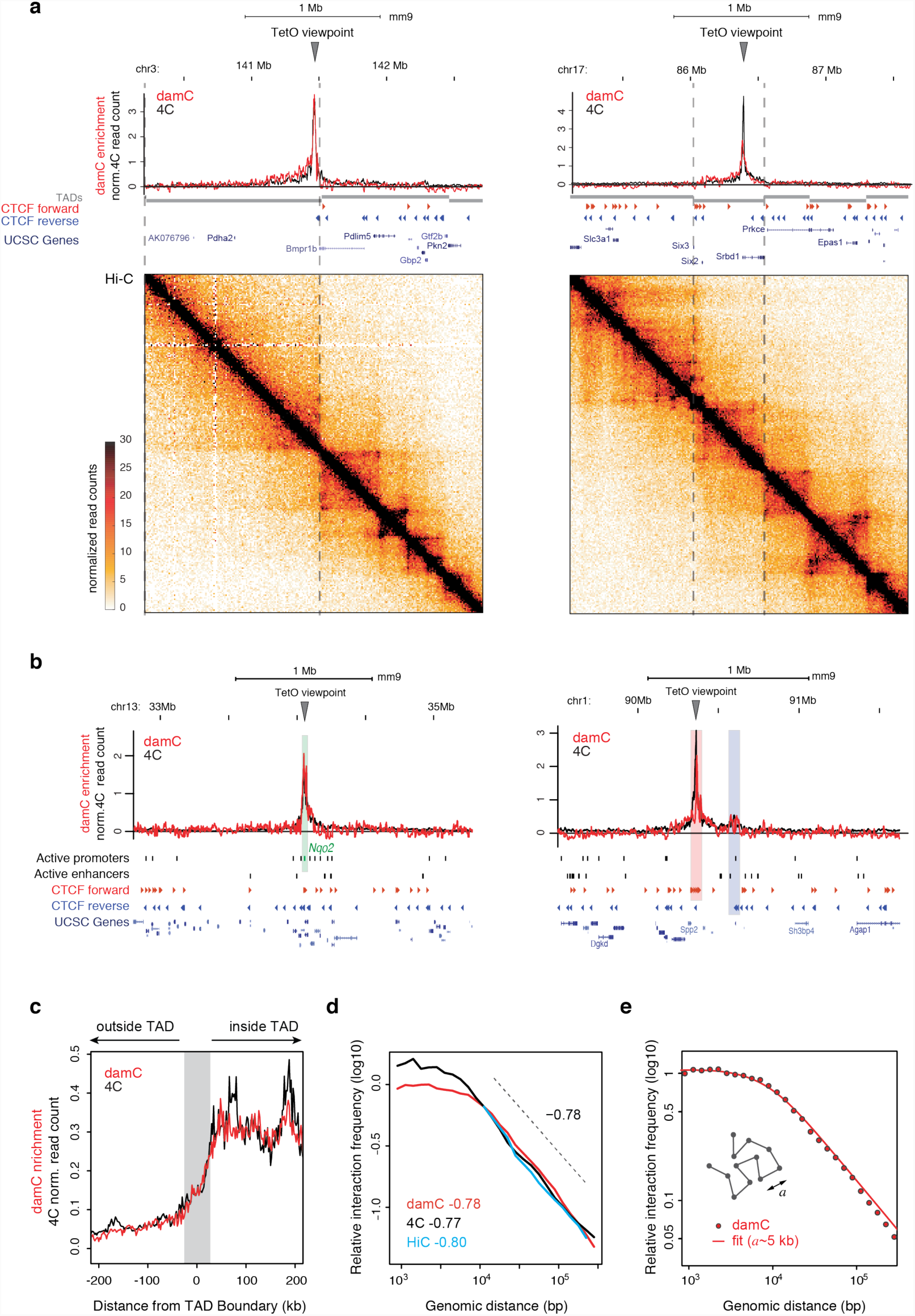
damC confirms the existence of TAD boundaries and quantitatively correlates with 4C and Hi-C. **a)** Two representative damC and 4C interaction from the same piggyBac-TetO viewpoints, aligned with Hi-C experiments performed in the same cell line. Dashed lines mark TAD boundaries in mESC detected using CaTCH^9^. Hi-C data were binned at 10 kb resolution. damC was performed using 10 nM 4-OHT. **b)** Interaction profiles from viewpoints located <1kb from the active promoter of the *Nqo2* gene (left), highlighted in the green shaded area, and <1kb from a CTCF site (right) belonging to a cluster of forward sites (red shading) interacting with reverse CTCF sites (blue shading). **c)** Aggregated plot over 100 TetO viewpoints showing damC and 4C data aligned to TAD boundaries identified using CaTCH^9^. Gray shading: +/-40 kb uncertainty on boundary definition^9^. **d)** Scaling of contact probabilities measured in damC, 4C and Hi-C from the same 100 viewpoints. **e)** Best fit of scaling measured by damC with a polymer model with persistence length *a.* The best value of *a* extracted from the fit is ∼5 kb.

We then asked whether damC and 4C measured the same scaling of interaction probabilities. We pooled all viewpoints and plotted the data as a function of genomic distance from the TetO viewpoints. For distances larger than ∼6 kb, fitting both damC and 4C with a power law resulted in a decay exponent around 0.78, in excellent agreement with Hi-C from the same cells and the same viewpoints (**Figure 4d**), and in accordance with previous measurements in the same range of genomic distances^14^.

Below ∼10 kb, both damC and 4C showed a gentler decay as recently observed in Hi-C experiments on yeast chromosomes^37^. In Ref.^37^ this effect was attributed to crosslinking artefacts but the fact that damC shows the same behavior argues against this explanation. The leveling-off of contact probabilities at short genomic distances can be explained in terms of a simple coarse-grained polymer model with a persistence length of 5 kb (**Figure 4e, Supplementary Model Description**). Thus although we cannot formally rule out alternative explanations (such as experimental factors not described by the damC model and thus not normalized in the calculation of enrichment), damC measurements suggest that *in vivo*, mouse chromosomes are essentially rigid below 5 kb.

We next investigated whether despite the global similarities, damC and 4C showed any local differences. To this aim we defined a deviation score measuring the differences between single damC and 4C interaction profiles within windows of 20 DpnI/II restriction fragments, corresponding to approximately 5kb on average (**Supplementary Figure 3d**). The most dissimilar 20-fragment windows were enriched in active and poised (polycomb-associated) chromatin marks compared to the most similar windows (**Supplementary Figure 3e**). We found that this is largely due to the fact that in both -Dox and +Dox conditions, the methylation signal is highly correlated with chromatin accessibility measured by DNase sensitivity (**Supplementary Figure 3f**). In the damC enrichment, correction by nonspecific methylation generally normalizes for chromatin accessibility (**Figure 2c**). Nevertheless a small number (7%, see Methods) of GATC sites within DNase hypersensitive regions are already highly methylated in the absence of Dox, preventing further increases in methylation due to interactions with the TetO viewpoint in +Dox conditions (**Supplementary Figure 3g**). Masking DNase hypersensitive sites indeed increases the similarity between damC and 4C (**Supplementary Figure 3h**).

In summary, crosslinking-and ligation-free measurements of *in vivo* contact probabilities using damC show quantitative agreement with 4C, confirm the existence of TAD boundaries, fully support 4C and Hi-C-based measurements of scaling exponents within TADs, and suggest that mammalian chromosomes are flexible only above 5 kilobases.

### piggyBac-TetO insertions do not perturb chromosome structure

Next, we set off to understand if the introduction of the TetO/piggyBac cassettes resulted in local perturbations in chromosome structure compared to wild-type cells. To this aim we compared TetO insertion sites with the corresponding wild-type loci in Hi-C experiments binned at 10 kb resolution (**Supplementary Figure 4a**). We used a modified version of the deviation score defined in **Supplementary Figure 3d** describing differences in virtual 4C profiles extracted from Hi-C data within 200kb windows (**Supplementary Figure 4b**). Deviation scores between wild-type cells and cells carrying TetO arrays were similar to those obtained when comparing two Hi-C replicates at random genomic locations, and significantly smaller than those derived comparing different wild-type loci (**Supplementary Figure 4b**). Finally, 4C profiles obtained in the region shown in Figure 4b (right panel) in the presence and absence of the TetO viewpoint were indistinguishable (**Supplementary Figure 4c**), indicating that the presence of TetO arrays did not perturb endogenous CTCF loops. Thus, insertion of TetO arrays through piggyBac transposition does not lead to any measurable effect on local chromosome structure.

### *In vivo* detection and manipulation of CTCF-mediated interactions

Loops between convergently oriented CTCF sites have emerged as a defining feature of chromosome architecture. However, no evidence exists that new loops can be established between endogenous and ectopically inserted CTCF sites. Since piggyBac-TetO constructs alone do not perturb chromosome structure, we reasoned that we could further engineer them to insert ectopic CTCF sites in the genome and detect the resulting structural modifications without confounding effects.

Starting from the founder rTetR-GFP-Dam-ERT2 mESC line described in Figure 3a, we introduced random insertions of a modified piggyBac cassette where the TetO array is flanked by three CTCF sites oriented outwards (**Figure 5a**). We reasoned that once inserted in the genome, the ectopic CTCF sites might establish loops with endogenous CTCF sites in the genomic neighborhood of the insertion (**Figure 5b**). To test this hypothesis we selected one clone carrying 93 insertions of the TetO-CTCF viewpoints, for which we could map the insertion position and genomic orientation (**Supplementary Table S4**), and performed damC in the presence of 10 nM 4-OHT, as well as 4C from the same viewpoints.

**Figure 5.**
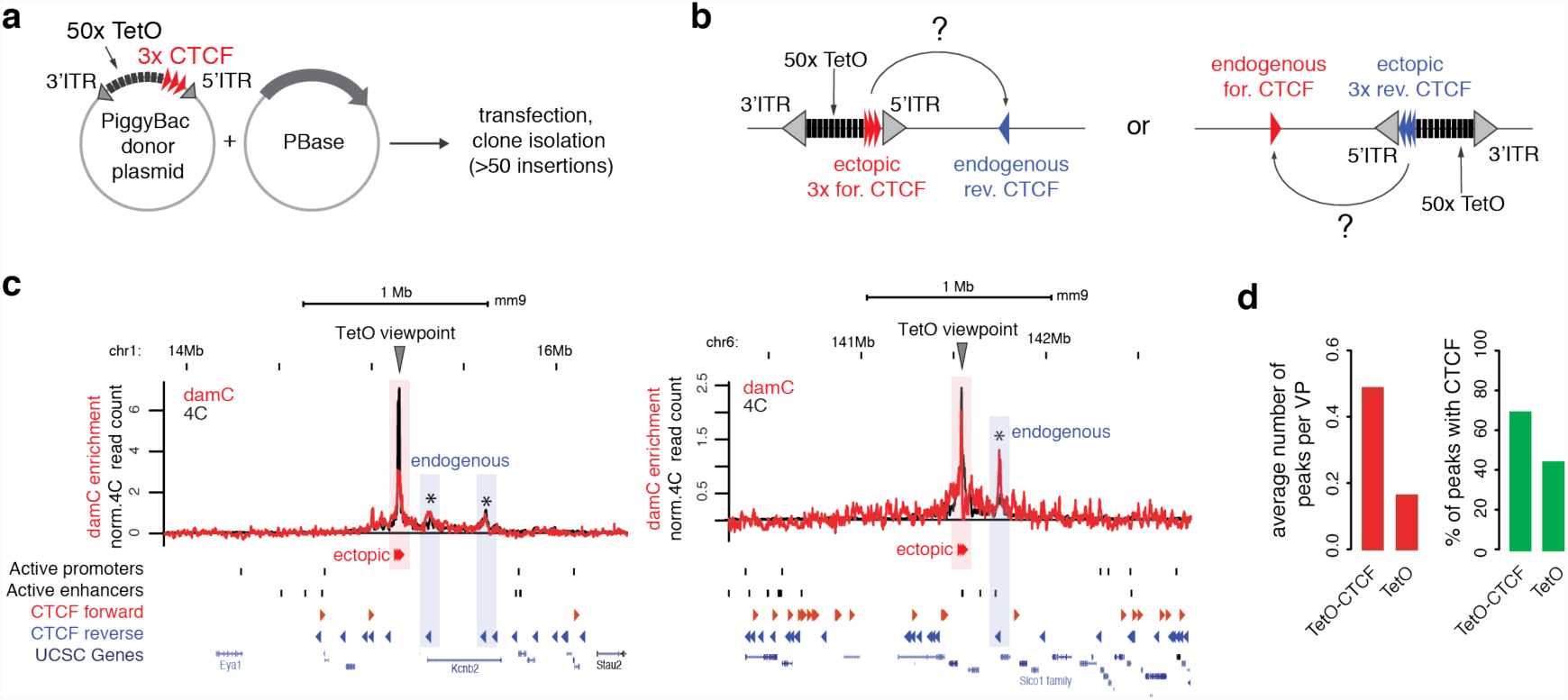
damC-based detection of CTCF loops. **a)** Modified piggyBac strategy to insert TetO viewpoints flanked by three CTCF sites oriented outwards. **b)** The TetO-CTCF cassette can insert in the genome in both directions and lead to the formation of interactions with either forward or reverse endogenous CTCF sites. **c)** Two representative interaction profiles obtained using damC and 4C from TetO-CTCF viewpoints. Asterisks indicate interactions identified by PeakC that overlap with CTCF sites. Shaded boxes indicate the overlap with the genomic positions of endogenous and ectopic CTCF locations. **d)** Left: average number of peaks per viewpoint detected by PeakC at least 20kb away from the viewpoint, in cells with TetO-CTCF or TetO-only insertions. Right: percentage of peaks containing a CTCF motif that is bound based on ChIP-exo data^11^.

Visual inspection of interaction profiles from single TetO-CTCF viewpoints revealed the presence of prominent distal peaks (**Figure 5c**), which are systematically detected in both damC and 4C. We used the PeakC algorithm, developed to analyze 4C profiles^38^, to systematically identify distal regions that preferentially interact with the TetO-CTCF viewpoints. Using stringent thresholds (see Methods section) we detected a total of 45 specific interactions separated by at least 20 kb from single TetO-CTCF viewpoints (around 0.5 distal peaks per insertion site on average). Of those, around 70% contained one or more bound CTCF sites based on ChIP-exo datasets in mouse ES cells^11^, in the vast majority of cases (77%) oriented in a convergent manner with respect to the ectopic CTCF insertion (**Figure 5d**). As a comparison in the cell line harboring TetO viewpoints without CTCF we detect only 0.16 peaks per insertion site, of which 43% contained one or more bound CTCF sites (**Figure 5d**). The fraction of viewpoints occurring in close proximity to an endogenous CTCF site is very similar in the TetO and TetO-CTCF insertion lines (**Supplementary Figure 5a**). Thus, peaks detected in the TetO-CTCF line are likely to coincide with new loops established as a consequence of the ectopic insertion of CTCF sites. Of the insertions without prominent distal interactions, the majority correspond to TetO-CTCF cassettes integrated relatively close (<30kb) to the nearest-neighbor convergent CTCF site, and/or in chromosomal regions that are very dense in endogenous CTCF sites, or CTCF deserts (**Supplementary Figure 5b**). This might lead to the formation of short-distance loops and structures that are more difficult to distinguish in the 4C and damC profiles.

To confirm that the CTCF-associated interactions we detected actually arise from new interactions that are absent in the wild-type locus, we performed Hi-C in the TetO-CTCF line and compared it to the data obtained from TetO-only mESCs (see **Figure 4a**), where insertion locations are different. It should be noted that TetO-CTCF insertions are heterozygous and therefore the Hi-C readout is confounded by the presence of a wild-type allele. Nevertheless, in a fraction of insertions showing prominent distal CTCF peaks in 4C and damC we could detect the formation of new structures in Hi-C and notably new loops (**Figure 6a** and **Supplementary Figure 5c**) leading to increased partitioning of interactions within TADs and the appearance of a sub-TAD boundary (**Figure 6b**). Interestingly, ectopic CTCF insertion did not only establish new loops with convergent endogenous CTCF sites (arrows in **Figure 6a**) but also reinforced pre-existing interactions between convergently oriented sites (arrowheads in **Figure 6a**), possibly by indirectly enforcing their proximity by effect of the new loops. Even insertions without prominent distal CTCF peaks could be associated to the formation of new structures (**Figure 6b**) reminiscent of stripes predicted by the loop extrusion model^16^ and recently observed in Hi-C data at endogenous locations^39^. Consistent with an interpretation in terms of loop extrusion by cohesin, the stripe-like feature shown in **Figure 6b** was induced at a location where the three ectopic CTCF sites landed close to a cluster of Nipbl sites, and far enough from the nearest convergent CTCF sites (∼800 kb) to allow the formation of a detectable stripe domain.

**Figure 6.**
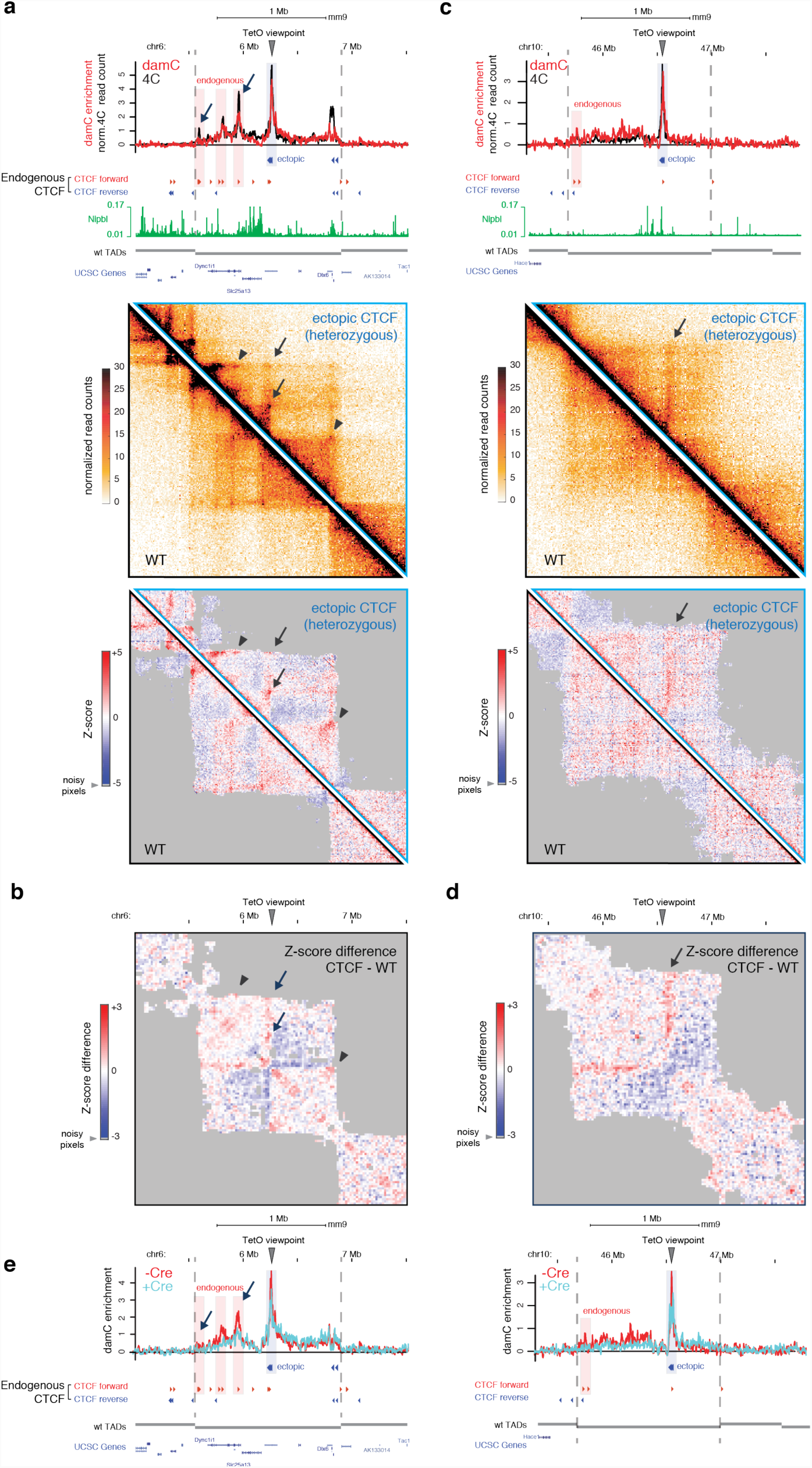
Ectopic CTCF insertion leads to the formation of new loops and stripes. a) TetO-CTCF insertion site giving rise to ectopic loops with convergently oriented endogenous CTCF sites. Top panel: Interaction profiles measured with damC and 4C are overlaid with the position of CTCF ChIP-exo sites from Ref.^11^ and Nipbl ChIP-seq data from Ref.^39^. Center panel: Hi-C data from the ESC lines carrying wither the heterozygous TetO-CTCF insertion or two wild-type alleles. Bottom panel: distance-normalized Z-scores, highlighting interactions that are either enriched (red) or depleted (blue) compared to the expected interaction frequency. Arrows: interactions between convergent CTCF sites that are established upon CTCF insertion. Arrowheads: pre-existing interaction that are strengthened after CTFC insertion. Hi-C data are binned at 10 kb resolution. **b)** Z-score difference between heterozygous CTCF and wild-type cells showing increased partitioning of interactions inside the TAD. Hi-C data were binned at 20kb. Shaded areas correspond to ‘noisy’ interactions that did not satisfy a quality control filter based on their correlations with immediate nearest neighbors (see Methods). **c)** Same as panel a for an insertion on chromosome 10, occurring in proximity to an isolated cluster of Nipbl binding and giving rise to a stripe-like interaction pattern. **d)** Z-score differences for the locus shown in panel c. **e)** damC interaction profiles from the same viewpoints as in panel a and c, before and after Cre-mediated excision of ectopically inserted CTCF sites (but not of the piggyBac cassette).

Formation of an ectopic CTCF-associated stripe also resulted in a modification of the partitioning of chromosomal interactions inside the TAD (**Figure 6d**).

Finally, to formally prove that the structures formed in the TetO-CTCF line are due to the presence of the three ectopic CTCF binding sites (rather than the piggyBac inverted terminal repeats or the TetO array), we removed the three CTCF sites by Cre-assisted recombination using two LoxP sites flanking the CTCF cassette (**Supplementary Figure 5d**). damC performed in one mESC clone where CTCF cassette had been excised at both loci shown in Figure 6a-b (**Supplementary Figure 5e**) revealed that interactions formed upon the ectopic insertion of CTCF sites were lost upon the removal of CTCF sites (**Figure 6e**).

In summary, damC readily identifies chromatin loops formed through specific long-range chromatin interactions. Additionally, our data shows that ectopically inserted CTCF sites can establish new structures (including loops and stripes) by interacting with endogenous CTCF sites that can lead to modified partitioning of interactions within TADs.

## Discussion

In this work we provide the first in vivo, high resolution, systematic measurements of chromatin contacts that do not require crosslinking nor ligation. Three independent studies previously employed the targeting of bacterial DNA adenine methyltransferase Dam into specific DNA sequences to identify spatial proximity in yeast, Drosophila and mammalian cells^25,31,32^. Each study however only involved the targeting of a single locus and used a semi-quantitative read-out to measure interactions. Here we move beyond these qualitative single-locus observations and systematically demonstrate at hundreds of genomic sites that damC can be used for the in vivo analysis of chromatin interaction frequencies. Notably, the use of next-generation sequencing to analyze contact frequencies in an unbiased manner allows appreciating these frequencies in a quantitative way and in the context of contacts made by all other surrounding genomic sequences. This allowed us to independently confirm the existence of CTCF-associated loops and TADs *in vivo*.

Unlike previous DamID-based approaches and in contrast to 3C-based methods, an essential feature of damC is that its experimental output is directly proportional to contact probabilities with the TetO viewpoints. This is supported by rigorous physical modeling of methylation kinetics (**Figure 2**), providing a rational basis for the quantitative interpretation of sequencing results. Importantly, damC confirms that contact frequencies drop across TAD boundaries by approximately a factor 2, in accordance with 4C (**Figure 4c**) and previous estimations based on Hi-C^16,40^. Such modest decrease raises the question of how TAD boundaries can achieve to functionally insulate enhancers and promoters from a purely biophysical point of view. However this relatively small decay in contact probabilities might represent the best compromise between enriching physical interactions between regulatory regions within topological domains and depleting them across boundaries^9^.

It should be noted that damC detects chromosomal contacts on very short spatial distances, since the adenine in a GATC motif can only be methylated if Dam is directly bound to DNA. We estimate the detection range to be smaller than 10nm, given that the expected physical size of the rTetR-EGFP-Dam-ERT2 fusion protein does not exceed 3 nm^41^. Decreases in interaction frequencies at TAD boundaries, as well as increases due to CTCF loops, therefore closely match what a promoter would ‘detect’ through protein complexes bound to its regulatory sequences. In this context it is also interesting to notice that damC picks up ‘non-specific’ interactions due to random collisions within the chromatin fiber to the same extent as 4C and Hi-C do (see for example **Figure 4a** and **6a**). Thus, random collisions do occur *in vivo*, despite not being detected in crosslinking-(but not ligation-) free analysis of chromosome folding using native 3C^25^.

Another remarkable finding is that the scaling of contact probabilities measured by damC is the same as those measured using 4C and Hi-C (**Figure 4d**). This is important because scaling behaviors extracted from Hi-C data are at the very core of virtually all physical models that have been proposed to date to explain chromosome folding and infer its mechanistic determinants^16,42–45^, including the highly influential loop extrusion model. Since the damC enrichment is proportional to chromosomal contact probabilities, our measurements provide strong evidence in favor of chromosome folding models based on polymer physics and Hi-C data. In addition, scaling analysis of damC data at short genomic distances in terms of a simple semiflexible polymer model suggests that mouse chromosomes might have a persistence length of approximately 5 kb, corresponding to ∼80 nm assuming the linear density of ∼60 bp/nm estimated in Ref.^42^.

The finding that loops can be established *de novo* upon insertion of CTCF binding sites, and can be detected *in vivo* (**Figure 5** and **6**) provides further evidence in favor of the involvement of CTCF in creating specific chromosomal interactions. These results also show for the first time to our knowledge that it is possible to manipulate chromosome structure in a ‘gain of function’ manner, by adding new structures rather than modifying existing ones. Notably, new structures formed upon the ectopic insertion of three CTCF sites can significantly modify the partitioning of intra-TAD interactions. At least a fraction of them results in the formation of new boundaries within pre-existing TADs, which can be detected in Hi-C despite the fact that piggyBac insertions are heterozygous (**Figure 6b** and **6d**). Remarkably, we could only detect newly formed interactions within pre-existing TAD boundaries. This is possibly a consequence of the fact that TAD boundaries are particularly enriched in clusters of CTCF binding sites^7,9^, providing efficient barriers to loop extrusion.

Finally, damC can readily be adapted to detect chromosomal interactions in a tissue-specific context by expressing the rTetR-Dam fusion under a tissue-specific promoter^46^, and starting from small numbers of cells^47^. Thus an exciting perspective will be to apply damC to study chromosomal interactions in situations where the cell number is a fundamental limiting factor (e.g. single embryos or rare adult cell types), and where 4C cannot be used. Targeted recruitment strategies such as CRISPR/dCas9 will additionally allow the interrogation of interactions of selected promoters or enhancers.

In summary, damC is the first crosslinking-and ligation-free method for the quantitative analysis of chromatin interaction frequencies. Besides providing an orthogonal validation of 3C-based findings and demonstrating that site-specific chromosomal interactions can be *de novo* established through the ectopic insertion of CTCF sites, damC opens up the possibility to detect chromosomal interactions *in vivo*, using small numbers of cells and in a tissue-specific way.

## Supplementary Figures

**Supplementary Figure 1.**
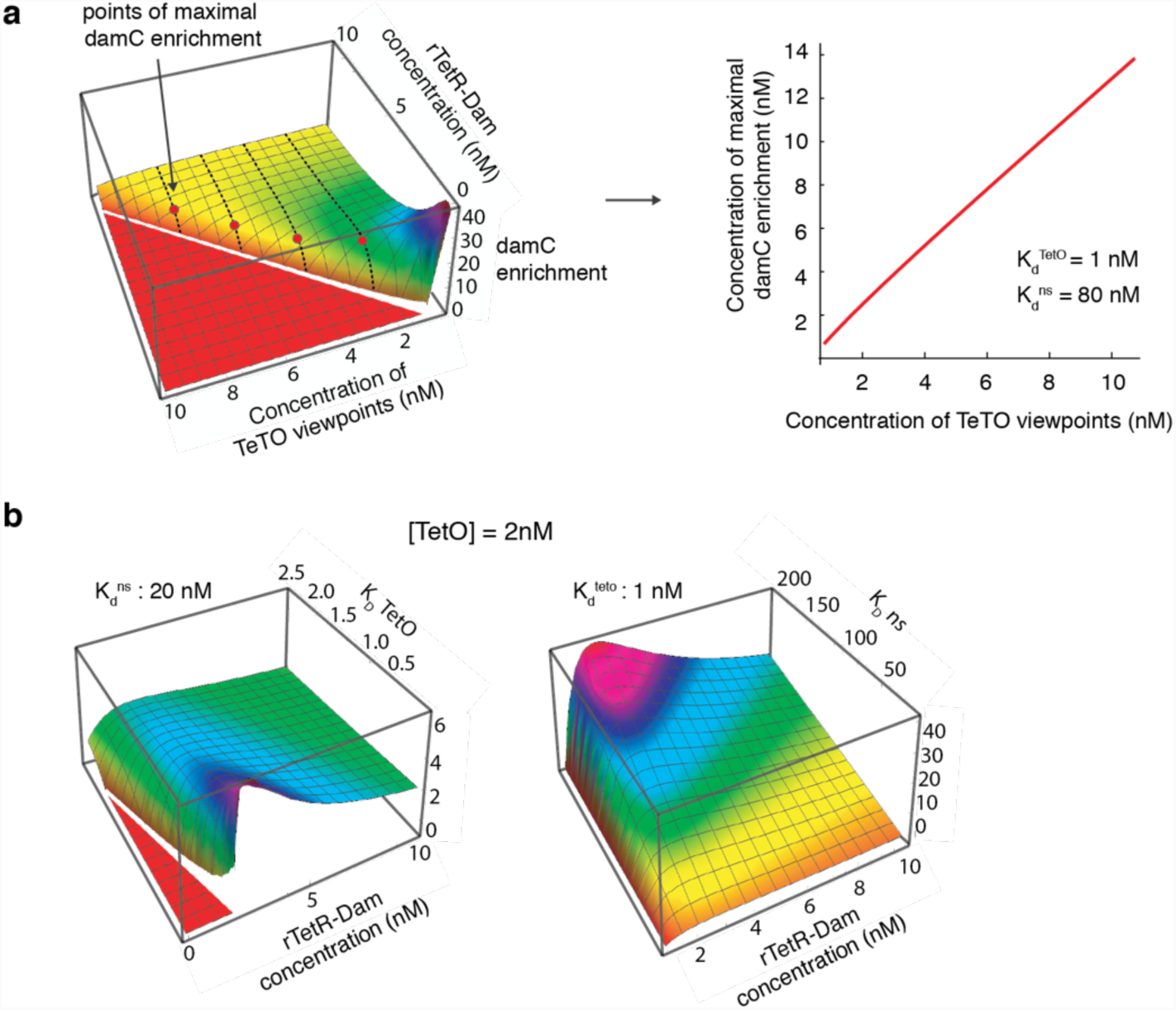
**a)** Left: damC enrichment is plotted as a function of the concentrations of rTetR-Dam and TetO viewpoints, imposing specific and non-specific dissociation constants to 1 nM and 80 nM respectively. Right: the rTetR-Dam concentration where the damC enrichment is maximal is linearly correlated with the concentration of TetO viewpoint. **b)** damC enrichment shows a maximum irrespective of the choice of the numerical parameters. This is exemplified by plots of damC enrichment as a function of rTetR-Dam concentration when varying the TetO specific affinity and keeping the nonspecific affinity fixed (left panel) and vice versa (right panel).

**Supplementary Figure 2.**
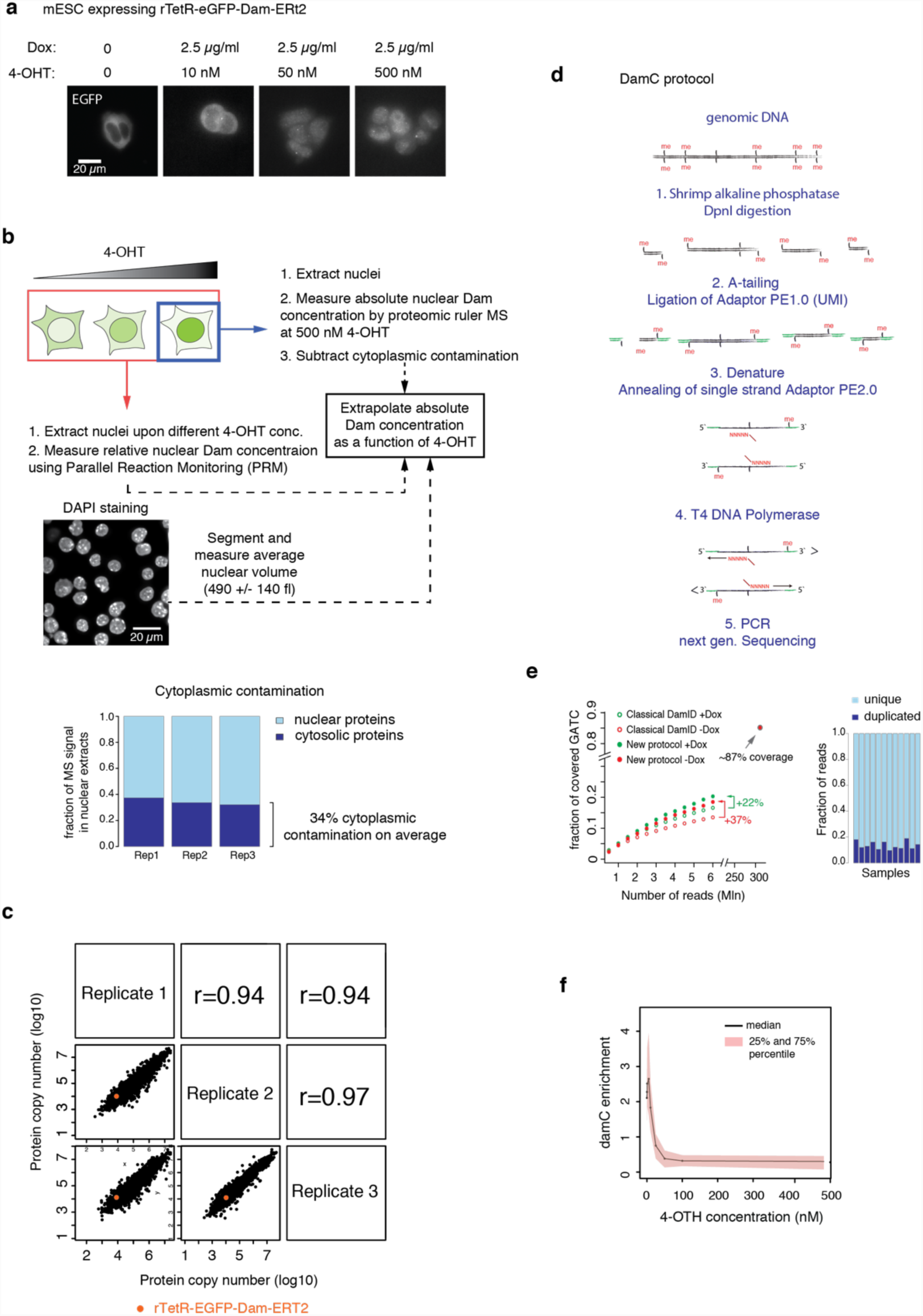
**a)** rTetR-Dam-EGFP-ERT2 becomes increasingly localized to the nucleus upon increasing 4-OHT concentration in the culture medium, as shown by the increasingly nuclear accumulation of EGFP. Maximum intensity projections of 10 wide-field Z planes are shown. Bright spots indicate binding of rTetR-Dam-EGFP-ERT2 to the 256x TetO array on chromosome X (see Figure 1c). **b)** Schematics of the strategy for measuring rTetR-Dam-EGFP-ERT2 nuclear concentrations as a function of 4-OHT concentration. After exposing the cells to different concentrations of 4-OHT, nuclei were extracted and prepared for mass spectrometry. The relative abundance of nuclear rTetR-EGFP-Dam-ERT2 was measured using parallel reaction monitoring (PRM) using two replicate samples from all 4-OHT concentrations. Absolute quantification was performed in triplicate uniquely in the 500 nM 4-OHT sample using proteomic-ruler mass spectrometry^34^. We then extrapolated absolute nuclear rTetR-Dam copy numbers at all concentrations of 4-OHT based on the absolute quantification at 500 nM 4-OHT and the relative PRM quantification. Finally, the nuclear concentration of Dam-fusion Protein was calculated based on the average nuclear volume determined based on DAPI staining. Contamination from cytoplasmic proteins was estimated by comparing protein copy numbers of nuclear and whole-cell extracts, and subtracted from nuclear copy numbers. **c)** Protein copy numbers determined in nuclear extracts at 500 nM 4-OHT using the proteomic ruler strategy34. Data from three biological replicates are plotted before correction for cytoplasmic contamination. **d)** Schematics of the damC library preparation. Genomic DNA is extracted from cells expressing the Dam-fusion protein. To avoid nonspecific ligation events in step 2, DNA is treated with shrimp alkaline phosphatase prior to DpnI digestion. After digestion with DpnI, a non-templated adenine is added to the 3’ blunt end of double-stranded DNA followed by ligation of the UMI-Adapter. Next, double-stranded DNA is denatured before random annealing of the second single stranded Adapter. In step 4, a T4-DNA-Polymerase is used for removal of 3’ overhangs and synthesis in the 5´ → 3´ direction. Finally, libraries are amplified by PCR and prepared for next generation sequencing. UMI: Unique Molecular Identifier. **e)** The damC sequencing library preparation protocol includes UMIs allowing to filter ∼15% of duplicated reads, and increases by roughly 30% the coverage of methylated GATC sites genome-wide compared to classical DamID30 at the same sequencing depth. **f)** Median DamC enrichment at 100 viewpoints with highest enrichment as a function of 4-OHT concentration. Significant amounts of damC enrichment can be observed in a range of rTetR-Dam nuclear concentrations corresponding to 5-10 nM 4-OHT in our experimental system.

**Supplementary Figure 3.**
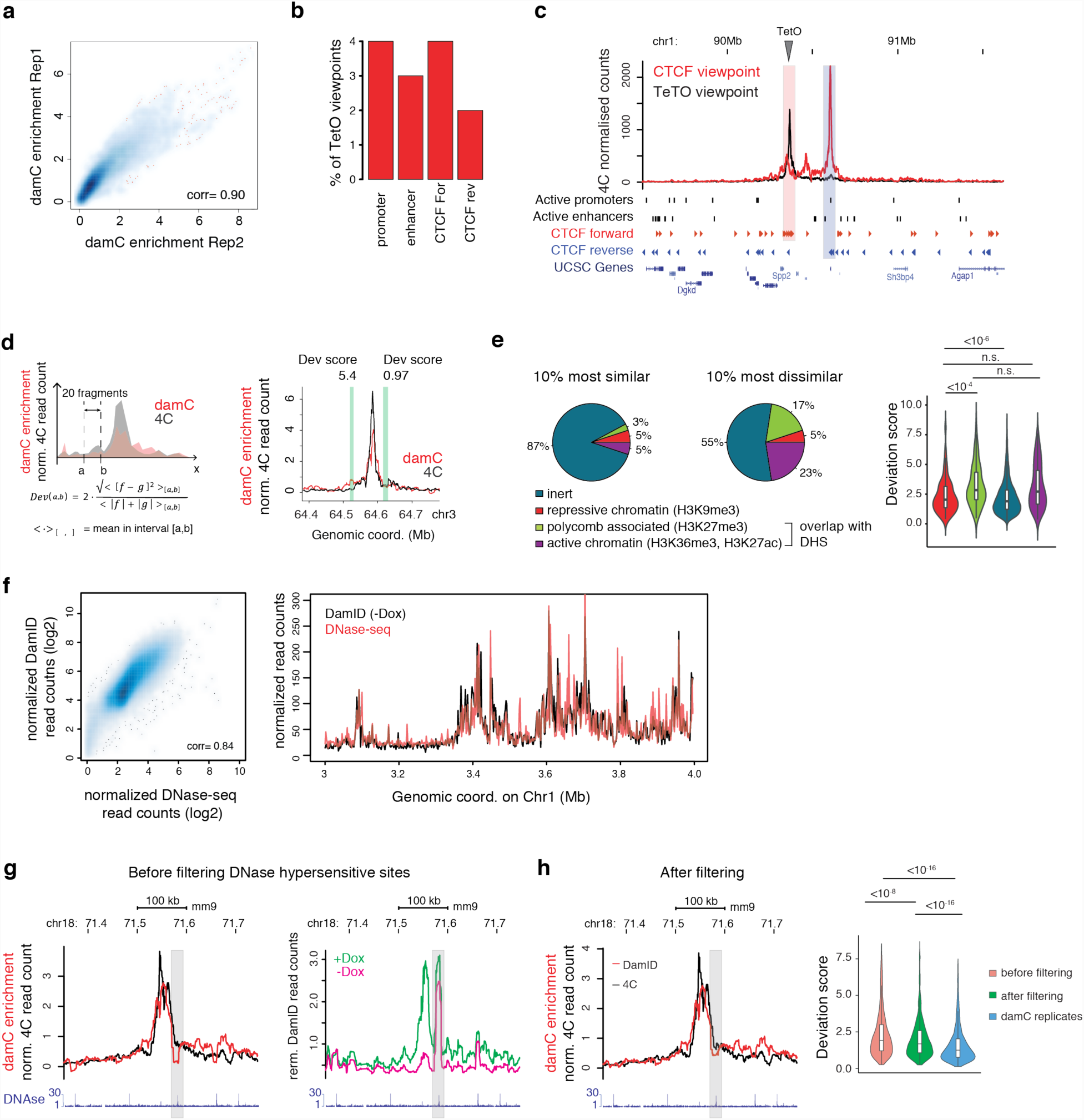
**a)** damC enrichment from single DpnI fragments within +/-100 kb from individual TetO viewpoints is plotted for two biological replicates performed with 10 nM 4-OHT. The Spearman correlation coefficient between the two replicates is indicated. **b)** The percentage of TetO viewpoints inserted in close proximity (<1 kb) from an active promoter or enhancer, or from a CTCF site that is bound in ChIP-exo^11^. **c)** 4C interaction profiles obtained at the same location as in the right panel of main Figure 3b, using the TetO viewpoint and the partner CTCF locus as a reverse viewpoint. **d)** Definition of a deviation score measuring local differences between damC and 4C. The deviation score is defined as the average quadratic difference between the damC and the 4C signal in a 20-restriction fragment interval, normalized by the mean of the signal in the same interval. Two intervals are shown on the right to illustrate the differences between deviation scores of ∼1 and ∼5. **e)** Right: the 10% most dissimilar 20-fragment intervals are enriched in active and bivalent (polycomb-associated chromatin) chromatin, based on the dominant ChromHMM state^48^ in the interval using four chromatin states (ChromHMM emissions) (Chi-Square Test: pvalue < 10-12). ‘Inert’ corresponds to chromatin that is not enriched in H3K9me3, H3K27m3, H3K36me3, H3K9ac, nor H3K27ac. See the Methods section for more details. Right: The distributions of deviation scores in 20-fragment intervals where the dominant ChromHMM state is either inert, repressive, polycomb-associated or active, showing that active or bivalent chromatin tends to show higher local dissimilarity between 4C and damC (pvalues from Wilcoxon test, two-sided). **f)** Left: correlation between damC signal in the -Dox sample and DNase-seq in mESC from ENCODE datasets. Each point in the scatter plot represents the aggregated signal in 20 kb; all 20kb intervals genome-wide are shown along with their Spearman correlation. Right: One representative megabase on Chr1 showing the high correlation between the two signals. damC and DNase-seq data were normalized to have equal average signal over the genomic interval shown here. **g)** Left: Representative damC interaction profile with GATC sites overlapping a DNase hypersensitive site that show saturated methylation levels (gray shaded area). Right: Normalized damC read counts in the same genomic region in the +Dox and -Dox samples highlight high basal methylation and limited increase in methylation, respectively, in the shaded area. **h)** Left: Removing DNase hypersensitive GATCs (see Methods) leads to increased local similarity between damC and 4C within the region plotted in panel g. Right: Distributions of local deviation scores calculated over all profiles, showing increased similarity between damC and 4C after removing DNase hypersensitive sites. Deviation scores between two damC biological replicates is shown for comparison (pvalues from Wilcoxon test, two-sided).

**Supplementary Figure 4.**
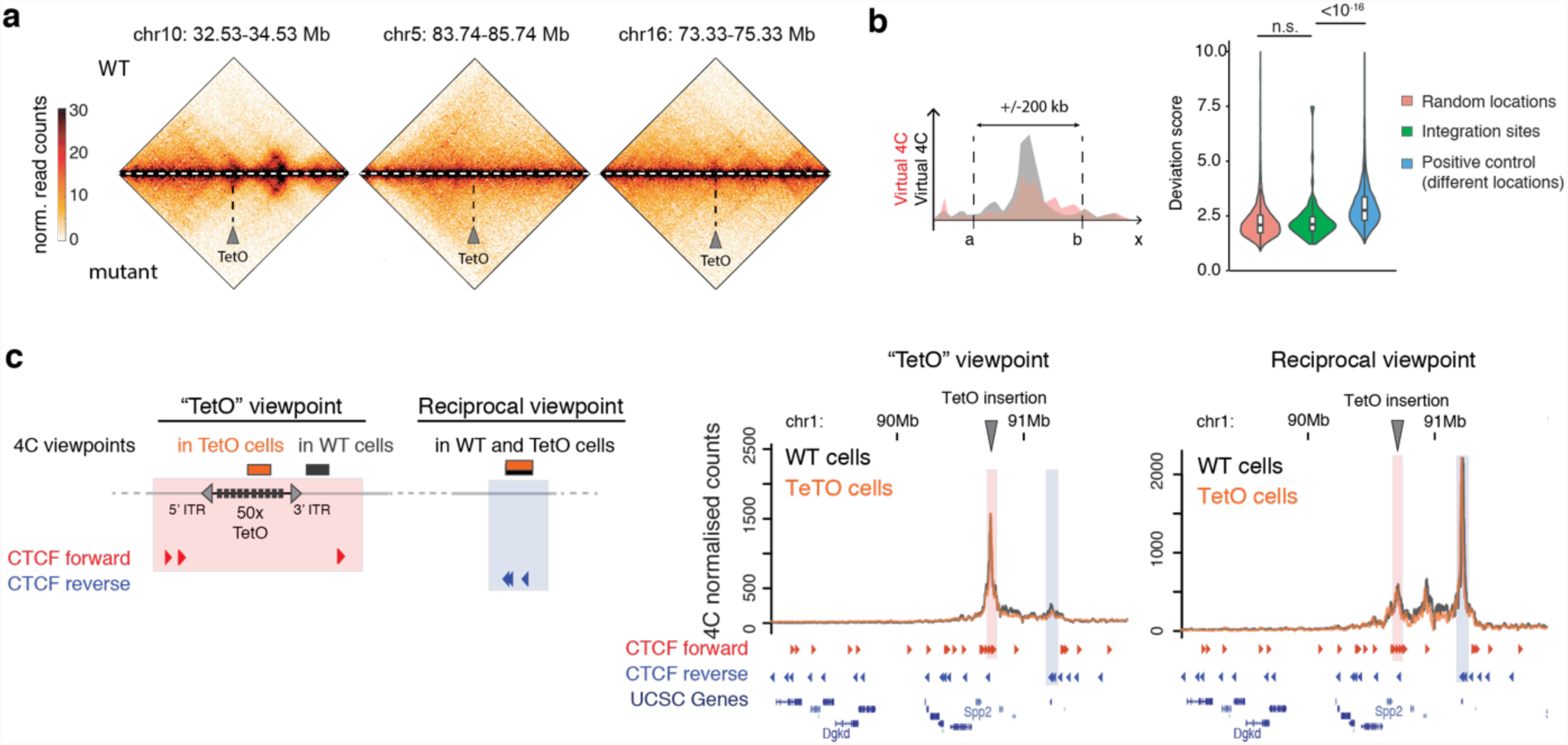
**a)** Insertion of TetO arrays does not perturb genome structure. Hi-C heatmaps of three different genomic locations harboring an array of 50xTetO sites and the corresponding wild-type locus are shown. Hi-C data are binned at 10 kb resolution. **b)** In a window of +/-200 kb surrounding the TetO integration sites, no significant changes can be detected in Hi-C at 10 kb resolution. Indeed, deviation scores (left) obtained at insertion sites (right, green violin plot) are similar to those obtained from random, non-insertion genomic viewpoints (pink violin plot), and significantly smaller than those obtained by comparing virtual 4C profiles from pairs of *different* random genomic viewpoints (blue) (pvalues are from Wilcoxon test, two-sided). **c)** Left: scheme of viewpoints used for the 4C experiment shown on the right. In cells harboring the TetO insertions, the ‘forward’ 4C viewpoint is within the TetO array as in main Figure 3; in wild-type cells, the viewpoint is adjacent to the insertion genomic coordinate. The reciprocal viewpoint is the same in the two cases. Right: 4C profiles at the locus shown in panel c using the viewpoints shown on the left are indistinguishable.

**Supplementary Figure 5.**
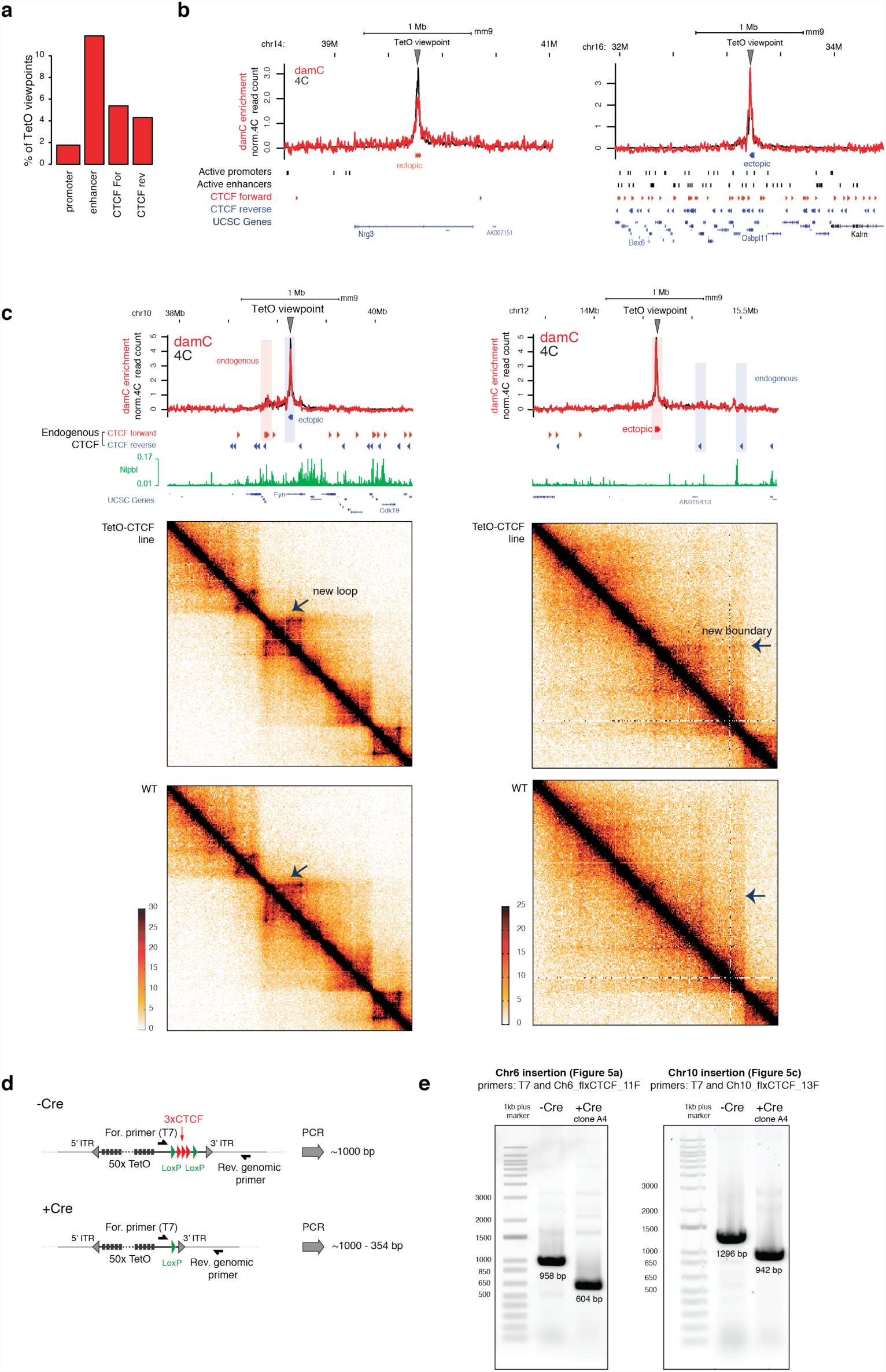
**a)** Percentage of TetO-CTCF viewpoints occurring in close proximity (<1 kb) from an active promoter or enhancer, or a CTCF site that is bound in ChIP-exo^11^. **b)** Examples of interaction profiles from TetO-CTCF viewpoints occurring in regions that are either devoid of (left) or densely bound by CTCF (right). **c)** Two further examples of ectopic structures formed as a consequence of the insertion of TetO-CTCF viewpoints. Hi-C data are binned at 10 kb resolution. **d)** Scheme of Cre-mediated excision of the ectopic CTCF cassette and genotyping. **e)** Genotyping PCR showing Cre-mediated excision of the CTCF cassette from the two integration sites shown in Figure 5 in the same mESC clone (A4).

## METHODS

### Physical modeling

Detailed descriptions of the physical model of methylation kinetics in damC, as well as of the polymer model with persistence length are available as a separate file (**Supplementary Model Description**).

### Cell culture and sample collection

All cell lines are based on feeder-independent PGK12.1 female mouse embryonic stem cells (mESC), kindly provided by Edith Heard’s laboratory. The founder cell line in our study is an X0 sub-clone of the PGKT2 clone described in (Masui et. al. 2011), carrying the insertion of a 256x TetO array within the 3’UTR of the *Chic1* gene on chromosome X and the additional deletion of the *Linx* promoter^6^ Cells were cultured on gelatin-coated culture plates in Dulbecco Modified Eagle’s medium (Sigma) in the presence of 15% foetal calf serum (Eurobio Abcys), 100 µM β-mercaptoethanol, 20 U/ml leukemia inhibitory factor (Miltenyi Biotec, premium grade) in 8% CO2 at 37°C. After insertion of the rTetR-Dam vector (see below), cells were cultured in the presence of 250 µg/mL hygromycin. To induce nuclear translocation of the rTetR-Dam fusion protein to the nuclei, mESC were trypsinized and directly seeded in culture medium containing 4-hydroxy-tamoxifen (4-OHT) at the concentrations indicated in the main text for 18 hours. Binding of the Dam fusion protein to the TetO arrays was induced by simultaneously adding 2.5 μg/ml doxycycline (Dox).

### Generation of cell lines expressing rTetR-Dam and carrying random insertions of TetO arrays

The rTetR-EGFP-Dam-ERt2 construct was cloned into a pBroad3 backbone (Invivogen) carrying a mouse Rosa26 promoter. We used a modified rTetR based on the rtTA-M2 transactivator in Ref.^49^, which has substantially decreased affinity for the Tet operator in the absence of Dox. The construct was randomly integrated in the PGKT2 X0 subclone by co-transfecting 5×10^5^ cells with 3 μg pBROAD3-rTetR-ICP22-EGFP-EcoDam-Ert2 and 0.2 μg of pcDNA3.1hygro plasmid using Lipofectamin 2000 (Thermo Fisher Scientific). After 10 days of hygromycin selection (250 μg/ml), one clone (#94.1) expressing low levels of EGFP was selected and expanded for subsequent experiments. To obtain large numbers of viewpoints for damC experiments, stable random integrations of arrays of Tet operator (TetO) sites were introduced in the #94.1 mESC clone using the piggyBac transposon system. A mouse codon optimized version of the piggyBac transposase^35^ was cloned in frame with the red fluorescent protein tagRFPt (Evrogen) into a pBroad3 vector using Gibson assembly cloning (pBroad3_hyPBase_IRES_tagRFPt). 5×10^5^ cells were co-transfected with 0.2ug of pBroad3_hyPBase_IRES_tagRFPt and 1µg of a piggyBac donor vector containing an array of 50 TetO binding sites using Lipofectamin 2000 (Thermo Fisher Scientific). Cells with high levels of RFP were FACS sorted two days after transfection and seeded at three serial 10x dilutions in 10-cm dishes to ensure optimal density for colony picking. To identify clones with high numbers of TetO integration sites, cells were screened for large numbers of nuclear EGFP accumulation foci using live-cell imaging (see below) in the presence of 500nM 4-OHT and 2.5ug/ml Dox and one clone (#94.1_2.7) was further expanded.

To introduce CTCF binding sites flanking the TetO viewpoints, the piggyBac donor vector was modified as follows. Three CTCF binding motifs (TGGCCAGCAGGGGGCGCTG, CGGCCAGCAGGTGGCGCCA and CGACCACCAGGGGGCGCTG) were selected based on high CTCF occupancy in ChIP-exo experiments^11^ and cloned into the piggyBac donor vector in an outwards direction with respect to the TetO array, including 100bp of their surrounding endogenous genomic sequence (chr8:13461990-13462089, chr1:34275307-34275419 and chr4:132806684-132806807, respectively). The three CTCF binding motifs were flanked by two LoxP sites for CRE assisted recombination. 5×10^5^ #94.1 were co-transfected with 0.2ug of pBroad3_hyPBase_IRES_rfp and 1µg of the modified piggyBac donor vector using Lipofectamin 2000. Cells with high levels of RFP were FACS sorted two days after transfection and seeded at three serial 10x dilutions in 10-cm dishes for colony picking. Clones with >50 of integration sites were identified through accumulation of EGFP at nuclear TetO foci in the presence of 500nM 4-OHT and 2.5ug/ml dox. One clone (#94.1_216_C3) was further selected for analyis.

### Transient transfection

To transiently express rTetR-Dam for the proof-of-principle experiment in Figure 1d, the PKGT2 X0 subclone was transiently transfected with pBroad3-rTetR-EGFP-Dam-ERt2 using the Amaxa 4D-Nucleofector X-Unit and the P3 Primary Cell 4D-Nucleofector X Kit (Lonza). 5×10^6^ cells were resuspended in 100 µl transfection solution (82ul primary solution, 18ul supplement 1, 2μg pBroad3-rTetR-EGFP-Dam-ERt2) and transferred in a single Nucleocuvette (Lonza). Nucleofection was done using the protocol CG109. Transfected cells were directly seeded in pre-warmed 37°C culture medium containing 10nM 4-OHT +/-2.5 μg/ml Dox. Genomic DNA was collected 18 hours after transfection. Sequencing libraries were prepared as previously described^30,50^.

### Mapping of piggyBac insertion sites

2µg of genomic DNA were fragmented to an average of 500bp by sonication (Covaris S220, duty cycle: 5%, peak power: 175W, duration: 25sec). End-repair, A-tailing and ligation of full-length barcoded Illumina adapters were performed using the TruSeq DNA PCR-free kit (Illumina) according to the manufacturer guidelines with the exception that large DNA fragments were not removed. 750ng of libraries for each sample were pooled together, and fragments of interest were captured using biotinylated probes against the the piggyBac inverted terminal repeats (ITRs) sequence and the xGen Hybridisation Capture kit (IDT) according to the manufacturer protocol (probes concentration of 2.25pmol/µl). Following the capture, libraries were amplified for 12 cycles using the Kapa Hi-fi polymerase and the following primers: 5’-AATGATACGGCGACCACCGAGAT, 5’-CAAGCAGAAGACGGCATACGAGA. Final libraries were purified using AMPure XP beads (1:1 ratio), quality controlled and sequenced on the NextSeq500 platform (paired-end 300 cycles mid-output) for a total of 8×10^8^ paired-end reads per sample on average.

. Capture probe sequences are as follows: ITR3-1 [Btn]ATCTATAACAAGAAAATATATATATAATAAGTTATCACGTAAGTAGAACATGAAATAACAATATAATTATCGTATGAGTTAAATCTTAAAAGTCACGTAAAAGATAATCATGCGTCATTT, ITR3-2 [Btn]TCCAAGCGGCGACTGAGATGTCCTAAATGCACAGCGACGGATTCGCGCTATTTAGAAAGAGAGAGCAATATTTCAAGAATGCATGCGTCAATTTTACGCAGACTATCTTTCTAGGGTTAA, ITR5-1 [Btn]TTAACCCTAGAAAGATAATCATATTGTGACGTACGTTAAAGATAATCATGCGTAAAATTGACGCATGTGTTTTATCGGTCTGTATATCGAGGTTTATTTATTAATTTGAA, ITR5-2 [Btn]ATTAAGTTTTATTATATTTACACTTACATACTAATAATAAATTCAACAAACAATTTATTTATGTTTAT TTATTTATTAAAAAAAAACAAAA ACTCAAAATTTCTTCTATAAAGTAACAAA.

### Genotyping of CTCF integration sites by PCR

We designed primers binding to endogenous genomic DNA sequence outside the piggyBac 3’ ITR based on the genomic position of mapped piggyBac insertion sites. We then amplified the junction between the ITR and the genome using Phusion High-Fidelity DNA Polymerase (Thermo Scientific) with one genomic primer and a T7 promoter primer (5’TAATACGACTCACTATAGGG3’) flanking the piggyBac CTCF integration cassette (see **Supplementary Figure 4d**). PCR products were purified and Sanger sequenced. For the verification of CTCF integrations shown in **Figure 4** and **Supplementary Figure 4e** on chromosome 6 and chromosome 10, the following genomic primers were used: Ch6_flxCTCF_11F (5’AGGCATTCTGTCCAACTGGT3’) and Chr10_flxCTCF_13F (5’TGTTGAGCATCTATCACATTCCTTA3’).

### Excision of CTCF sites using Cre recombinase

In order to excise ectopically inserted CTCF sites from the #94.1_216_C3 clone, 5×10^5^ cells were transfected with 0.5 μg of pIC-Cre^51^ using Lipofectamine 2000 (Thermo Fisher Scientific). After 4 days under G418 selection (300 ug/ml), single colonies were expanded and genotyped following the procedure described above.

### Live-cell Imaging

Gridded-glass-bottom dishes (Mattek) were coated with 2 ug/ml recombinant mouse E-cadherin (R&D Systems, 748-EC) in PBS at 4°C overnight. 5×10^5^ cells were seeded in full medium one day before imaging supplemented with 4-OHT and Dox as indicated above. Cells were imaged with a Nikon Eclipse Ti-E inverted widefield microscope (Perfect Focus System with real time drift correction for live cell imaging) operating in TIRF mode using a CFI APO TIRF 100x/1.49 oil objective (Nikon). A 488nm, 200mW Toptica iBEAM SMART laser was used as excitation source. Cells were maintained at a constant temperature of 37°C and 8% CO2 within an incubation box. Images were collected with an Evolve(tm) 512 Delta EMCCD high speed Camerang using Visiview (Visitron). Background subtraction (150-pixel rolling ball radius) and maximum intensity projections were performed in ImageJ.

### Nuclear volume measurements

3×10^6^ Cells from the #94.1_2.7 mESC clone were cultured in gelatin-coated 6-well plates in full medium, dissociated 5 minutes at room temperature with Accutase (GIBCO), then centrifuged 4 minutes at 950 rpm and resuspended in 500 µl of culture medium. 25-µl droplets of cell suspension were spotted on coverslips previously coated with poly-L-lysine, let adsorb on ice for 5 minutes and washed gently once with 1X PBS. Cells were then permeabilized on ice for 5 min in 1X PBS and 0.5%Triton X-100, and coverslips were stored in 70% EtOH at −20C. Nuclei were counterstained with 0.2 mg/ml DAPI and Z-stack images were acquired using a Zeiss Z-1 microscope equipped with a 40x oil immersion (NA=1.3) (voxel size 0.227×0.227×0.73 µm). Z-stacks were then deconvolved using the Huygens software (20 iterations of the CMLE algorithm). To segment individual nuclei, we binarized DAPI images based on a single intensity threshold based on the fact that image histograms of all Z-stacks were bimodal (threshold = 7000 in 32-bit images). The volumes of binary 3D objects was then calculated using the 3DObjectCounter plugin in FIJI/ImageJ, excluding objects on the edges of each Z-stack.

### Preparation of nuclear extracts

Cell nuclei were extracted as previously described^52^. Briefly, 10^7^ mES cells were seeded in ES medium (see above) supplemented with the appropriate concentration of 4-OHT on a gelatin coated 15 cm^2^ dish. The next day, cells were harvested using trypsin and washed twice in ice cold PBS. Next, cells were carefully resuspended in 500μl ice cold Buffer A1 (10mM HEPES pH7.9, 10mM KCl, 1.5mM MgCl2, 0.34M sucrose, 10% glycerol, 0.1% Triton-X 100, 1mM DTT, 1mM PMSF) to obtain nuclei. After incubating for 5 minutes on ice, extracted nuclei were washed twice with buffer A1.

### Mass spectrometry

Nuclear extracts were dissolved in 400 µL 50mM HEPES pH 8.5 in 8.3M guanidine hydrochloride. All the samples were heated at 95°C for 5 min, sonicated using Bioruptor^®^ sonication device and supplemented with 5mM TCEP and 10mM CAA. To reduce sample complexity, lysates were diluted to 6M guanidine hydrochloride and transferred onto 100kDa molecular weight cut-off Amiconultra-0.5 centrifugal filter units. Samples were concentrated for 2 x 15 minutes at 14kG followed by refill of the filter with 6M guanidine hydrochloride in 50mM HEPES pH 8.5 and 3 x 45 minutes at 14kG followed by refill of the filter with 1M guanidine hydrochloride in 50mM HEPES pH 8.5. For digestion 10µg of Lys-C (Wako Chemicals) and 10µg of trypsin (Thermo Fisher) were added to each sample and incubated over night at 37°C. In the morning additional 10µg of trypsin was added, incubated for 3h and acidified using TFA.

To estimate nuclear copy numbers samples were desalted using SEP-PAK (Waters) and subjected to high pH offline fractionation on a YMC Triart C18 0.5 x250 mm column (YMC Europe GmbH) using the Agilent 1100 system (Agilent Technologies). 96 fractions were collected for each experiment and concatenated into 48 fractions as previously described^53^. For each LC-MS analysis, approximately 1 µg of peptides were loaded onto a PepMap 100 C18 2 cm trap (Thermo Fisher) using the Proxeon NanoLC-1000 system (Thermo Fisher). On-line peptide separation was performed on the 15 cm EASY-Spray C18 column (ES801, Thermo Fisher) by applying a linear gradient of increasing ACN concentration at a flow rate of 150 nL/min. An Orbitrap Fusion Tribrid (Thermo Fisher) or an Orbitrap Fusion LUMOS Tribrid (Thermo Fisher) mass spectrometers were operated in a data-dependent mode. The top 10 most intense precursor ions from the Orbitrap survey scan were selected for higher-energy collisional dissociation fragmentation and analyzed using the ion-trap.

### Mass spectrometry data processing

Maxquant version 1.5.3.8 was used to search raw mass spectrometry data using default settings^54^ against the mouse protein sequences from Uniprot database (release 2017-04). The label free quantification (LFQ)^55^ algorithm was used for quantification. The protein groups table were loaded to Perseus software^56^ (version 1.5.0.0) filtered for potential contaminants and reverse hits. Protein copy numbers per cell were calculated using the Protein ruler plugin of Perseus by standardization to the total histone MS signal^34^ (Wisniewski et al., 2014). The LFQ values were normalized using same normalization for all samples. To estimate cytoplasmic contamination “GOCC slim name” annotations provided in Perseus were used. Exclusively cytoplasmic proteins were defined as being associated with GOCC terms “cytoplasm” or “cytosol” and not associated with terms “nucleus”, “nuclear”, “nucleoplasm” and “nucleosome”. Exclusively nuclear proteins were defined as being associated with GOCC terms “nucleus”, “nuclear”, “nucleoplasm” and “nucleosome” and not associated with terms “cytoplasm” or “cytosol”. The cytoplasmic contamination was estimated using a ratio of summed LFQ intensity between exclusively cytoplasmic proteins and exclusively nuclear proteins in samples with and without nuclear extraction.

### Parallel reaction monitoring (PRM) data acquisition and analysis

To select peptides for PRM assays, the rTetR-Dam-EGFP-ERT2 construct was enriched using magnetic ChromoTek’s GFP-Trap beads and analyzed using shotgun data-dependent acquisition LC-MS/MS on an Orbitrap Fusion Lumos platform as decribed above. For PRM analysis the resolution of the orbitrap was set to 240k FWHM (at 200 m/z), the fill time was set to 1000 ms and ion isolation window was set to 0.7 Th. The acquired PRM data were processed using Skyline 4.135^57^. The transition selection was systematically verified and adjusted when necessary to ensure that no co-eluting contaminant distorted quantification based on traces co-elution (retention time) and the correlation between the relative intensities of the endogenous fragment ion traces, and their counterparts from the library.

### damC library preparation

DamC experiments are based on a newly developed DamID-seq NGS library preparation protocol to maximize the proportionality between methylation levels and sequencing readout (**Supplementary Figure 2c**). One crucial issue in the calculation of enrichment as in **Figure 2c** is that small fluctuations in -Dox methylation in the denominator can be amplified into large fluctuations in enrichment levels. GATC sites must therefore be equally and robustly represented in the DamID sequencing library irrespective of their methylation level. From this perspective, the principal limitation of the original DamID protocol^30^ for our present application was its dependence on the genomic distance between two GmATC sites, resulting in large adaptor-ligated molecules and as a consequence in a strong bias towards densely methylated regions. In our optimized protocol, GmATC sites are sequenced independently of the neighboring GATC methylation status resulting in a ∼30% increase in GmATC coverage at equivalent sequencing/read depth (**Supplementary Figure 2d**). In addition, we introduced unique molecular identifiers (UMIs) allowing a precise enrichment quantification after excluding PCR duplicates from the sequencing data.

Overall, the damC library construction protocol can be divided in 3 parts: 1) ligation of UMI adapters with a “one-tube” strategy, 2) integration of the second sequencing adapter, followed by 3) a final PCR amplification. Briefly, 3×10^6^ cells were harvested using trypsin after 18 hours of induction with tamoxifen +/-doxycyclin. Genomic DNA was extracted using the Qiagen blood and tissue kit adding 250U of RNaseA in step 1. Genomic DNA was eluted in 80ul ddH2O. DNA concentration was measured using the Qbit DNA Broad Range kit. Genomic DNA (100ng input) was treated with Shrimp Alkaline Phosphatase treatment (NEB, 1U), followed by DpnI digestion (ThermoFisher Scientific, 10U), A-tailing (0.6mM final dATP, 5U Klenow exo-, ThermoFisher Scientific), and UMI adapters ligation (30U T4 DNA ligase, PEG4000, ThermoFisher Scientific) performed within the same tube and buffer (Tango 1X, ThermoFisher Scientific) by heat inactivating each enzymatic step followed by adjustment with the reagents required for the next step. UMI adapters were made by annealing the following oligos: 5’-AATGATACGGCGACCACCGAGATCTACACNNNNNNNNACACTCTTTCCCTACACGACGCTCTTCCGA TC*T and 5’-pGATCGGAAGAGCGTCGTGTAGGGAAAGAGTGT. Ligation reactions were treated with Exonuclease I (20U, ThermoFisher Scientific) then purified using AMPureXP beads (1:0.8 ratio, Agencourt) and the second sequencing adapter (5’ TGACTGGAGTTCAGACGTGTGCTCTTCCGATCTNNNNN*N 3’, IDT) was tagged using heat denaturation and second strand synthesis (5U T4 DNA Polymerase, ThermoFisher Scientific). The tagging reaction was purified using AMPure XP beads (1:1 ratio) followed by a final library amplification (10 cycles) using 1U of Phusion polymerase, 2μl 10 μM DAM_UMIindex_PCR (5’ AATGATACGGCGACCACCGAGATCTACA*C 3’), and 2μl 10μM NEBnext indexed primer (NEB). Final libraries were purified AMPure XP beads (1:1 ratio) and QCed using Bioanalyser and Qbit. DamC libraries were sequenced on a NextSeq500 (75 cycles single-end) with a custom sequencing protocol (dark cycles at the start of read1 to “skip” the remaining DpnI site TC sequence). Samples index were determined using index1 read, and UMI sequence using index2 read. We obtained the following read counts: #94.1_2.7 cell line (TetO insertions only), 1.5×10^8^ valid reads on average per sample (+/-Dox, two replicates each). For the #94.1_216_C3 line (TetO-CTCF), 3.8×10^7^ valid reads on average per sample; for the Cre-excised clone A4, 4.3×10^7^ reads per sample. Detailed number of total and valid reads can be found in **Supplementary Table S6**.

### 4C-seq

4C sample preparation was performed as previously described^58^. Briefly, 10^7^ cells were cross-linked in 2% formaldehyde for 10 minutes and quenched with glycine (final concentration 0.125M). Cells were lysed in 150 mM NaCl/50 mM Tris-HCl (pH 7.5)/5 mM EDTA/0.5% NP-40/1% Triton X-100. The first digest was performed with 200 U DpnII (NEB), followed by ligation at 16 ° C with 50 U T4 DNA ligase (Roche) in 7 mL. Ligated samples were de-crosslinked with Proteinase K (0.05 ug/uL) at 65° C, purified, and digested with 50 U Csp6I (Thermo Fisher Scientific) each, followed by ligation with 100 U T4 DNA ligase in 14 mL and purification. Resulting products were directly used as PCR template for genomic dedicated 4C viewpoints. Primers for PCR were designed using guidelines described previously^58^. We obtained the following read counts: #94.1_2.7 cell line (TetO insertions only), 4.2×10^6^ valid reads on average per sample (+Dox, two replicates). For the #94.1_216_C3 line (TetO-CTCF), 3.5×10^6^ valid reads on average per sample; for the experiments shown in Supplementary Figure 3c and 4c, we obtained an average of 3.2×10^6^ reads per sample. Detailed number of total and valid reads can be found in **Supplementary Table S6**.

### In vitro Cas9 digestion of 4C templates

In order to detect chromosomal interactions directly from the genome integrated TetO platform, viewpoint primers were designed to amplify directly from the DpnII fragments contained in the TetO sequence. The 2.7 kb TetO platform contains a total of 50x contiguous repeats of the same TetO DpnI/II viewpoint. To prevent PCR amplification and sequencing of TetO repeats due to tandem ligation of two or more TetO DpnII fragments in a given 4C circle, an in vitro Cas9 digestion was performed on the 4C templates. Cas9 was targeted into the TetO repeats in between viewpoint primers using a single gRNA. In vitro transcribed gRNA template was obtained using the Megashortscript T7 transcription kit (Invitrogen). gRNA was purified with 4 × AMPure purification (Agencourt). Purified Cas9 protein was kindly provided by N. Geijsen. Cas9 was pre-incubated with the sgRNA for 30 min at 37 °C. Subsequently, 4C template DNA was added to the pre-incubated gRNA-Cas9 complex and incubated for 3–6 h at 37 °C for digestion. Cas9 was inactivated by incubating at 70 °C for 5 min.

### Hi-C library preparation

6×10^6^ mESC were harvested and diluted in 1x PBS to final 1×10^6^ cells/ml, then crosslinked with 1% formaldehyde and quenched with 0,125M glycine for 5 min at RT. After two 1x PBS washes, cells pellets were obtained by centrifugation, snap frozen and stored at −80°C. Pellets were thawed on ice and resuspended in 500 ul lysis buffer (10 nM Tris-HCl pH8.0, 10 nM NaCl, 0.2%NP40, 1x Roche protease inhibitors) and left 30 min on ice. Cells were then pelleted by centrifugation (954 x g, 5 min, 4°C), washed once with 300 ul 1x NEB2 buffer and nuclei were extracted by 1 h incubation at 37°C in 190 ul 0.5%SDS 1xNEB2 buffer. SDS was neutralized by diluting the sample with 400 ul NEB2 buffer and adding 10% Triton X-100. After 15 min of incubation at 37°C, nuclei were pelleted, washed once in PBS and resuspended in 300 ul NEB2 buffer. 400U of MboI (NEB, 25 000 units/ml) were added and incubated at 37°C overnight. The next day, nuclei were pelleted again, resuspended in 200 ul fresh NEB2 buffer and additional 200U of MboI were added for two more hours before heat inactivation at 65°C for 15 min. 43 ul of end-repair mix (1.5 μL of 10 mM dCTP; 1.5 μL of 10mM dGTP; 1.5 μL of 10 mM dTTP; 37.5 μL of 0.4 mM Biotin-11-dATP (Invitrogen) and 1 μL of 50U/μL DNA Polymerase I Large Klenow fragment (NEB)) were added to the nuclear suspension, incubated at 37°C for 45 min and heat inactivated at 65°C for 15 min. The end repair mix was exchanged with 1.2 ml of ligation mix (120μL of 10X T4 DNA Ligase Buffer; 100 μL of 10% Triton X-100; 6 μL of 20 mg/mL BSA; 969μL of H2O) plus 5 ul of T4 ligase (NEB, 2000 units/ml) and ligation was performed at 16°C overnight. Nuclei were reconstituted in 200 ul fresh NEB2 buffer followed by RNA digestion in 0.5 mg/ml RNAse A for 10 min at 37°C. Samples were de-crosslinked with Proteinase K at 65°C overnight and DNA was purified using phenol/chloroform. 2 ug of DNA sample were sonicated using Diagenode Bioruptor Pico. MyOne Streptavidin T1 (Life Technologies # 65601) magnetic beads were used to capture biotinylated DNA followed by A-tailing. Adapter ligation was performed according to NEB Next Ultra DNA Library prep kit instructions. Two independent PCR reactions with multiplex oligos for Illumina sequencing were performed and pooled for the final PCR clean-up by magnetic AMPure bead (Beckman Coulter) purification. The final libraries were eluted in nuclease-free water, QCed by Bioanalyzer and Qubit. HiC libraries were sequenced on a Illumina Nextseq500 platform (2×42bp paired end). We obtained an average of 2.9×10^8^ valid reads per sample (TetO-only and TetO-CTCF cells, -Dox, two biological replicates each). Detailed number of total and valid reads can be found in **Supplementary Table S6**.

### Sequencing data processing and data analysis

#### DamC analysis

All samples were aligned to mouse mm9 using qAlign (QuasR package^59^) using default parameters. PCR duplicates were removed using a custom script. Briefly, reads were considered PCR duplicates if they map to the same genomic location and have the same 8-bp UMI sequence. We quantified the number of reads mapped to each GATC that could be uniquely mapped using qCount (QuasR package^59^). The *query* object we used in qCount was a GRanges object containing the uniquely mappable 76-mers GATC loci in the genome shifted upstream (plus strand) or downstream (minus strand) by 5 base-pairs (3 dark cycles + *GA*, see Supplementary Figure 2c and the *‘*Bulk DamID-seq Library preparation’ paragraph in the Methods section). Each sample was then normalized to a common library size of 10M reads and a pseudo-count of 0.2 was added. Prior to calculating damC enrichments, a running average over 21 restriction fragments was performed and the mean value was assigned to the central GATC. Enrichment was then calculated as in Figure 2c: E=([+Dox]-[-Dox])/[-Dox] where [+Dox] and [-Dox] are the normalized and running-averaged number of reads in the presence and absence of Dox, respectively. We defined the damC signal to be saturated if it satisfies the following criteria: 1) it belongs to the top highest 25% genome wide both in +Dox and -Dox samples, and 2) the ratio between +Dox and -Dox methylation is close to 0.5, i.e. belongs to the [0.45, 0.55] quantile of all ratios genome wide.

#### 4C analysis

Mapping of 4C reads was performed as described for damC, with the exception of UMI de-duplication since 4C libraries did not include UMIs and quantification was done by counting the reads mapped exactly to the GATC sites. The two restriction fragments immediately flanking the piggyBac-TetO cassette were excluded from subsequent analyses.

#### Hi-C Analysis

Hi-C data were analysed using HiC-Pro version 2.7.10 with *--very-sensitive --end-to-end --reorder* option. Briefly, reads pairs were mapped to the mouse genome (build mm9). Chimeric reads were recovered after recognition of the ligation site. Only unique valid pairs were kept. Contact maps at a given binning size were then generated after dividing the genome into equally sized bins and applying iterative correction^61^ on binned data.

#### Fit of scaling plots

Average normalized Hi-C counts, damC enrichment or 4C counts were calculated for all pairs of loci separated by logarithmically binned distance intervals. The binning size in logarithmic scale (base 10) was 0.1. Curves were fitted in log-log scale using the lm function in R.

#### Fitting the damC model to damC experiments as a function of 4-OHT concentration

The damC enrichment depends on the rTetR-TetO specific and nonspecific dissociation constants, the concentration of TetO and the nuclear rTetR-Dam concentration (Supplementary Model Description). In addition, it depends on the actual contact probability between the genomic location where it is calculated and the TetO viewpoint. In Figure 3d we calculated the damC enrichment at the closest fragments to the 100 TetO viewpoint with higher signal to noise ratio. We assumed that the contact probability between the TetO array and the closest fragment is ∼1, and fitted the model to the experimental data using the other parameters with the *NonlinearModelFit* function in Mathematica. The constraints that the dissociation constants and the concentration of TetO are positive were imposed. The goodness of the fit was evaluated using the adjusted R^2^ (0.74).

#### ChromHMM

In order to assign chromatin states, we used the ChromHMM software^48^ with four states. We used histone modifications as in **Supplementary Table S5.** The four states correspond to active (enriched in H3K36me3, H3K27ac, H3K4me1 and H3K9ac), poised (H3K36me3, H3K27ac, H3K4me1, H3K9ac and H3K27me3), inert (no enrichment) and heterochromatic (H3K9me3) states.

#### Deviation scores

Given a set of restriction fragments (or genomic bins) {x_i_} belonging to a window [a,b], the deviation score is defined as

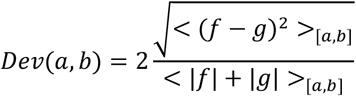

Where *f* and *g* are data vectors (e.g. damC enrichment, 4C or virtual 4C counts) and < >_[*a*,*b*]_represents the average in the window [a,b]. If two profiles are identical in the window [a,b], then the deviation score is zero; increasing deviation from zero indicates increasing dissimilarity.

#### PiggyBac-TetO integration site mapping

Paired-end reads (see ‘Mapping of piggyBac insertion sites’ above) were trimmed to 50 bp using a custom script. Read1 and Read2 were mapped separately to the piggyBac-TetO sequence using QuasR (qAlign). Only hybrid pairs with one of the reads mapping to array were kept. The second reads from hybrid pairs were mapped to the mouse genome (build mm9) using QuasR (qAlign). Reads were then piled up in 25 bp windows using csaw (windowCounts function). Integration sites can be identified because they correspond to local high read coverage. Local coverage was calculated by resizing all non-zero 25-bp windows up to 225 bp (expanding by 100 bp upstream and downstream). Overlapping windows were then merged using reduce (from GenomicRanges) thus resulting in a set of windows {*w*_*i*_}. The size distribution of *w*_*i*_ is multimodal, and only *w*_*i*_ from the second mode on were kept. For each *w*_*i*_ we estimated the coverage *c*_*i*_ as the number of non-zero 25-bp windows. Only *w*_*i*_’s where the coverage were higher than 16 were considered. The exact position of the integration sites were then identified with the center of *w*_*i*_.

#### Determination of the orientation of TetO-CTFC insertions

In order to determine the orientation of ectopically inserted TetO-CTCF sites, we exploited the fact that the three CTCF sites are oriented within the piggyBac casette in the 3’ ITR - 5’ ITR direction. If the genomic position of the 5’ ITR is upstream of the 3’ ITR, then CTCF sites are in the reverse orientation (-strand), and vice versa. To determine the relative orientation of the 3’ and 5’ ITRs in the genome, we used only reads that run through the junction between the ITRs and the genome. More precisely, we extracted reads that contain an exact match to 30bps of the ITRs (3’ and 5’ ITR separately), trimmed the ITR sequence and mapped the reads to the mouse genome using qAlign (from QuasR). We quantified the reads at single bp resolution using scanBam. Only integration sites where both 5’ and 3’ ITRs are mapped are kept. This resulted in 93 integration sites (**Supplementary Table S4**).

#### Z-score analysis of Hi-C data

In order to identify and exclude ‘noisy’ interactions in Hi-C maps we used a custom algorithm named ‘Neighborhood Coefficient of Variation’ (van Bemmel *et al*., under revision). Since the chromatin fiber behaves as a polymer, the contact probability of a given pair of genomic loci *i* and *j*, is correlate to that of fragments i+N and j+N if N is smaller (or on the order of) than the persistence length of the chromatin fiber. Hence, a given pixel in a Hi-C map can be defined as noisy if its numerical value is too different from those corresponding to neighboring interaction frequencies. To operatively assess the similarity between neighboring interactions, we calculated the coefficient of variation (CV) within a 10×10 pixel square centered on every interaction and discarded all pixels whose CV is larger than a certain threshold. Given that the distribution of the coefficient of variation of Hi-C samples in this study is multimodal with the first component terminating around CV=0.6, we set the CV threshold to 0.6. Discarded interactions appear as grey pixels in the differential Hi-C maps. For differential analysis between TetO-CTCF and wt samples, we calculated the difference between distance-normalized Z-scores calculated for each individual map^62^. The Z-score is defined as (obs-exp)/stdev where (obs) is the Hi-C signal for a given interaction and (exp) and (stdev) are the genome-wide average and standard deviation of Hi-C signals at the genomic distance separating the two loci.

#### 4C peak calling

In order to call specific interactions in 4C profiles, we used the peakC package^38^ using the following parameters: qWr = 2.5 and minDist = 20000. peakC was applied to 2 replicates of 4C profiles at single fragment resolution with 21 running average steps. Peak regions were then extended 1kb upstream and downstream. Overlapping peaks were merged.

## Acknowledgments

Research in the Giorgetti lab is funded by the Novartis Foundation and ERC Starting Grant n. 759366 ‘BioMeTRe’. We would like to thank Michael Stadler for help with bioinformatics analysis, Stefan Grzybek and Hans-Rudolf Hotz for assistance on server supports, Ralph Grand for help with CTCF site sequences, Edith Heard and Rafael Galupa for kindly providing PGK cells. We are grateful to Gioacchino Natoli, Dirk Schuebeler and Rafael Galupa for critically reading the manuscript. We acknowledge The ENCODE Project Consortium and in particular the Ren and Hardison laboratories for ChIP-Seq data sets in ESC. This work is dedicated to the memory of Maxime Dahan.

## Data availability

The sequencing data from this study have been submitted to the NCBI Gene Expression Omnibus (GEO; http://www.ncbi.nlm.nih.gov/geo).

## Author contributions

JR generated cell lines and performed damC experiments. YZ wrote the model with assistance from GT and analyzed the data. CV performed 4C in WdL’s lab. MK assisted with cell culture and damC library preparation and performed Hi-C experiments. IMG and JK helped with experimental design and data analysis. VI performed mass spectrometry experiments and analysis. TP provided constructs for initial experiments and discussed the data. SS developed the damC library preparation protocol and performed piggyBac insertion mapping experiments. LG designed the study and wrote the paper with JR and YZ and input from all the authors.

